# Transcript architecture predetermines m6A remodeling and sensory neuron vulnerability in chemotherapy-induced peripheral neuropathy

**DOI:** 10.64898/2026.05.18.722999

**Authors:** Md Mamunul Haque, Ohannes K Melemedjian

## Abstract

Whether individual transcripts carry intrinsic features that predetermine their response to perturbations is unknown. Here we used nanopore direct RNA sequencing of male mouse dorsal root ganglia (DRG) to simultaneously profile N6-methyladenosine (m6A) modifications, poly(A) tail dynamics, and full-length isoform identity from mice treated with bortezomib, a proteasome inhibitor that causes painful peripheral neuropathy. Machine learning revealed that transcript-intrinsic features predetermine the magnitude of perturbation-induced m6A loss (R² = 0.983). Expression level contributed just 2.6% of predictive importance. Bortezomib removed a fixed ∼73.5% fraction of m6A marks, meaning absolute loss scaled linearly with baseline density and a transcript’s epitranscriptomic fate was encoded in its architecture before drug exposure. Unsupervised clustering identified four response programs where the dominant m6A erosion cluster enriched for oxidative phosphorylation (OXPHOS, p = 1.0 × 10⁻¹⁷) and proteasome (p = 2.8 × 10⁻¹⁰) genes, recapitulating bortezomib’s established mechanisms without prior biological knowledge. Isoform-resolved analysis uncovered m6A remodeling patterns suggesting post-transcriptional regulation of glycolytic and OXPHOS genes, and Western blot confirmed protein-level suppression of OXPHOS components. Integration with single-nuclei sequencing showed sensory neurons carried 2.2-fold greater m6A loss burden than non-neuronal cells, a direct consequence of architectural determinism applied to cell-type-specific transcriptomes. These findings establish that epitranscriptomic bortezomib response is predetermined by transcript architecture, with pathway specificity and cell-type vulnerability emerging as downstream consequences of intrinsic RNA structure.

## INTRODUCTION

How cells respond to perturbations, whether pharmacological, pathological, or environmental, is conventionally understood through the lens of differential gene expression where a stimulus alters signaling pathways, transcription factors are activated or repressed, and mRNA levels change accordingly. This framework has dominated transcriptomics for two decades and has yielded important insights, but it rests on an assumption that changes in mRNA abundance are the primary readout of cellular response. Whether individual transcripts carry intrinsic architectural features that predetermine their molecular fate under perturbation, independent of transcriptional regulation, has not been systematically investigated.

This question cannot be addressed using conventional analytical platforms. Short-read or sequencing-by-synthesis (SBS) methods fragment RNA molecules, reverse-transcribe them into cDNA, and amplify them before sequencing. Each of these steps introduces biases and destroys information. Fragmentation eliminates single-molecule connectivity, reverse transcription erases native chemical modifications, and amplification distorts quantitative relationships[8; 22; 34]. The resulting data reduce each transcript to a single expression value, channeling analysis toward differential gene expression as the primary readout of biological change. These methods embed a blind spot because the information needed to detect regulatory events that alter protein output without changing transcript levels are destroyed before sequencing begins.

Nanopore direct RNA sequencing eliminates these biases by passing, native RNA molecules through a protein pore, recording base sequence, N6-methyladenosine (m6A) modifications, poly(A) tail length, and full-length isoform identity simultaneously from individual molecule[15; 26]. This single-molecule, multi-modal capability preserves the very information that short-read and SBS approaches destroy, enabling the measurement of regulatory layers that are invisible to conventional transcriptomics.

Here we exploit this capability to ask a fundamental question: do transcripts carry intrinsic architectural features that predetermine their response to perturbations? We use bortezomib-induced peripheral neuropathy (BIPN) as a model system because it offers three critical advantages. First, bortezomib, a proteasome inhibitor used in the treatment of multiple myeloma, induces painful peripheral neuropathy in up to 57% of treated patients [37], representing a severe drug-induced clinical pain condition with clear biological endpoints.

Second, the underlying biology is well characterized: bortezomib stabilizes HIF1α[16; 21], which upregulates enzymes such as lactate dehydrogenase A (LDHA), to drive a metabolic switch from oxidative phosphorylation to aerobic glycolysis in dorsal root ganglia (DRG)[16; 20]. This established mechanistic framework provides a stringent benchmark against which purely computational discoveries can be validated. Third, the post-transcriptional mechanisms underlying this metabolic reprogramming remain unknown, creating a clear opportunity for discovery.

By applying nanopore direct RNA sequencing to mouse DRG and employing machine learning, we discovered that transcript-intrinsic features, motif composition and spatial organization, predetermine the magnitude of perturbation-induced m6A loss, while expression contributes just 2.6% of predictive importance. Independent unsupervised analysis recapitulated bortezomib’s established mechanisms without prior biological knowledge. Isoform-resolved analyses uncovered m6A remodeling patterns suggesting post-transcriptional regulation of glycolytic and OXPHOS genes, with western blot confirming protein-level suppression of OXPHOS components. Integration with single-nuclei data demonstrated that sensory neuron vulnerability emerges directly from this architectural principle. These findings demonstrate that a transcript’s epitranscriptomic fate is written into its architecture before exposure occurs.

## METHODS

### Animal Model and Bortezomib Treatment

Pathogen-free, adult male ICR mice (3–4 weeks old; Envigo) were housed in a temperature (23°C ± 3°C) and light (12-h light/12-h dark cycle; lights on 07:00–19:00) controlled rooms with standard rodent chow and water available ad libitum. Animals were randomly assigned to treatment or control groups. All procedures were approved by Institutional Animal Care and Use Committee (IACUC protocol 0817005). Mice were treated with either intraperitoneal (IP) vehicle or 0.2 mg/kg of bortezomib (Millipore Sigma, Cat # 5.04314.0001) for five consecutive days for a total dose of 1 mg/kg. Dorsal root ganglia (DRG) were harvested at three timepoints: Control (4 groups, 20 mice total, n=5/group), Day 7 post-treatment (3 groups early CIPN, 15 mice total, n=5/group), and Day 14 post-treatment (3 groups established CIPN, 15 mice total, n=5/group). DRGs were rapidly dissected and frozen.

### RNA Extraction

The DRGs from 5 mice were pooled into each sample. Total RNA was isolated using QIAzol lysis reagent (Qiagen, Venlo, Netherlands, cat # 79306) according to the manufacturer’s instructions. The RNA was air dried for 10 minutes, resuspended in nuclease-free water, quantified, and either stored at −80°C or processed further by poly(A) purification. Dynabead mRNA purification kit (Thermo, cat # 61006) was used according to the manufacturer’s instructions to isolate poly(A) tail–containing mRNAs. The resulting poly(A) RNA was eluted in nuclease-free water and stored at −80°C.

### Nanopore Direct RNA Sequencing

Oxford MinION libraries were generated using the Direct RNA Sequencing Kit (Oxford Nanopore Technologies, Oxford, UK, cat # SQK-RNA002) according to the manufacturer’s instructions. Sequencing was done on MinION Mk1B (Oxford Nanopore Technologies) using R9.4.1 Flow Cells (Oxford Nanopore, cat # FLO-MIN106). A total of 10 libraries were sequenced: 4 Control replicates, 3 Day 7 replicates, and 3 Day 14 replicates.

### Basecalling and Alignment

Raw FAST5/POD5 signals were basecalled using Dorado 0.9.6. The –estimate-poly-a flag was enabled during basecalling to simultaneously estimate poly(A) tail lengths from the raw signal. Basecalled reads were aligned to the GRCm39 mouse reference genome using minimap2 with parameters optimized for direct RNA sequencing. Aligned reads were sorted and indexed using samtools.

### N6-methyladenosine (m6A) Detection

m6A modifications were detected at single-molecule resolution using m6Anet 2.1.0 [11], which applies a multiple-instance learning framework to identify m6A sites from nanopore current signal deviations. m6Anet was run on each of the 10 samples independently. The tool generates per-site modification probabilities (data.site_proba.csv) aggregated across reads, as well as per-read modification probabilities (data.indiv_proba.csv) that preserve single-molecule resolution. Per-read probabilities were mapped back to original read identifiers by indexing into the corresponding FASTQ files, enabling integration with poly(A) tail measurements on the same molecule.

### Poly(A) Tail Length Estimation

Poly(A) tail lengths were estimated directly from the raw nanopore signal during basecalling using Dorado’s built-in –estimate-poly-a algorithm, which identifies the poly(A) signal segment and converts its duration to nucleotide length. This approach provides per-read poly(A) measurements that are inherently matched to the same molecules carrying m6A modification data, enabling single-molecule multi-modal analysis without requiring separate computational tools.

### Isoform Discovery and Quantification

Full-length transcript isoforms were identified and quantified using FLAIR (Full-Length Alternative Isoform analysis of RNA) 3.0.0 [32]. The FLAIR pipeline was executed in the recommended unified collapse mode, where splice-site-corrected reads from all 10 samples were concatenated prior to isoform collapse, following the tool’s documentation for multi-sample experimental designs. The pipeline proceeded through the following steps: (1) alignment of reads to the GRCm39 genome, (2) splice-site correction against the GENCODE vM38 annotation, (3) concatenation of all corrected BED files across samples, (4) unified isoform collapse using the –stringent and –check_splice flags with read-to-isoform mapping enabled (–generate_map), (5) per-sample quantification against the unified isoform set using flair quantify with –quality 0, and (6) isoform productivity prediction using predictProductivity with the – longestORF flag, which classifies isoforms as productive (PRO), premature termination codon-containing (PTC), nonsense-mediated decay targets (NMD), no-go decay targets (NGO), or non-stop decay targets (NST).

### Multi-Modal Feature Matrix Construction Per-Site Feature Assembly

A comprehensive per-site feature matrix was constructed by integrating data from m6Anet, Dorado poly(A) estimates, and GENCODE vM38 transcript annotations. For each of the 10 samples, m6Anet per-read modification probabilities were merged with site-level kmer context (5-mer DRACH motif identity) and matched to Dorado poly(A) tail lengths via read identifiers. Transcript structural features including total exonic length, 5′ UTR, CDS, and 3′ UTR lengths were derived from the GENCODE vM38 GTF annotation. Positional features were calculated for each m6A site relative to the transcript body, including normalized position (0–1 scale), distance from transcript start and end, and regional classification (5′ UTR, CDS, or 3′ UTR based on 10%/90% positional thresholds).

### Sequencing Depth Normalization

To correct for inter-group differences in sequencing depth, normalization factors were calculated from the ratio of mapped reads per group relative to the Control condition. D7 samples exhibited 5.9% lower sequencing depth and D14 samples 1.0% lower depth compared to Controls.

Inverse correction factors of 1.063x for D7 and 1.010x for D14 (with Control as 1.000× reference) were applied multiplicatively to raw site and read counts prior to calculating any density-based metrics (e.g., m6A sites per kilobase), ensuring that observed differences in modification density reflected biological variation rather than technical library size effects.

### Transcript-Level Aggregation

Per-site observations were aggregated to transcript-level features within each sample, yielding one feature vector per transcript–sample combination. Features were organized into six categories: (1) m6A modification features (mean, median, and coefficient of variation of modification probability, percentage of high-confidence sites >0.7, m6A density as sites/kb); (2) poly(A) tail features (mean, median, standard deviation, coefficient of variation, interquartile range, skewness, kurtosis, median absolute deviation, and percentile values of poly(A) tail length distributions, plus the percentage of short (<50 nt) and long (>200 nt) tails); (3) structural/motif features (transcript length, counts of each of the 18 DRACH 5-mer motifs, motif diversity, and percentage of sites in DRACH context); (4) spatial features (mean and median normalized position, 3′ enrichment ratio, site counts by transcript quartile, position entropy, and spatial clustering index); (5) baseline features calculated from the mean of Control replicates for m6A and poly(A) metrics; and (6) temporal delta features representing the change from Control baseline for each treatment timepoint.

### Isoform Feature Integration

FLAIR-derived isoform features were merged with the transcript-level matrix at the gene level. For each gene, isoform complexity features were calculated from the FLAIR counts matrix, including the number of expressed isoforms, Shannon entropy of isoform usage proportions, dominant isoform fraction, and a binary isoform switching indicator (change of dominant isoform between conditions). Structural features included the mean, maximum, and minimum number of exons per isoform and isoform lengths. Isoform productivity features derived from FLAIR’s predictProductivity output included the number and percentage of productive isoforms, PTC-containing isoforms, NGO isoforms, NST isoforms, and NMD targets, as well as an overall coding potential flag. Isoform entropy changes between Control and treatment conditions (delta_isoform_entropy_d7, delta_isoform_entropy_d14) were computed as the difference in Shannon entropy of isoform usage proportions.

### Expression Data Integration

Gene-level expression values in transcripts per million (TPM) were obtained from the Oxford Nanopore wf-transcriptomes pipeline output and merged by gene name. Mean TPM values were calculated per condition (control_mean_tpm, d7_mean_tpm, d14_mean_tpm), and expression fold-changes were computed as log2((treatment_tpm + 1) / (control_tpm + 1)) with a pseudocount of 1 to handle zero-expression genes. The final feature matrix comprised 69 baseline/intrinsic features and 16 response variables. D7 and D14 timepoints were combined, and transcript–timepoint observations with complete data across all response variables were retained for downstream analysis.

### Machine Learning Prediction of Perturbation Response Data Preparation and Leakage Prevention

A strict data leakage prevention protocol was implemented to ensure that the predictive model used only features available prior to perturbation. The 69 predictor features were restricted to baseline measurements (derived from Control samples) and intrinsic transcript properties: 21 structural/motif features (transcript length, 18 DRACH motif counts, motif diversity, DRACH percentage), 9 spatial features (positional statistics and clustering metrics measured in Control), 6 baseline m6A features (sites/kb, modification probabilities, detection rate), 15 baseline poly(A) features (distributional statistics of tail lengths), 16 baseline isoform features (complexity, productivity, structure), and 2 baseline expression features (Control TPM). Treatment timepoint measurements and delta features were used exclusively as response variables, never as predictors.

### Response Variables

Sixteen response variables were defined to capture the multi-dimensional molecular response across four regulatory layers: 2 m6A responses (change in sites/kb from Control, change in mean modification probability), 11 poly(A) responses (changes in mean, median, coefficient of variation, interquartile range, median absolute deviation, skewness, and kurtosis of tail length; changes in percentage of short and long tails; and poly(A) CV ratio and mean ratio relative to baseline), 2 isoform responses (change in Shannon entropy, binary isoform switching), and 1 expression response (log2 fold-change). D7 and D14 timepoints were combined, and rows with any missing response variable values were excluded.

### Model Architecture and Training

A multi-output gradient boosting regression model was trained using scikit-learn’s MultiOutputRegressor wrapper around GradientBoostingRegressor. Each of the 16 response variables was modeled by an independent gradient boosting regressor sharing a common feature space. Hyperparameters were set as follows: 200 boosting iterations (n_estimators=200), maximum tree depth of 10, learning rate of 0.1, minimum samples for a split of 20, minimum samples per leaf of 10, and a fixed random seed of 42 for reproducibility. Features were standardized using z-score normalization (StandardScaler) prior to model fitting. Missing values in predictor features were imputed with the column median. The dataset was split into 80% training and 20% held-out test sets using random sampling (random_state=42). Model performance was evaluated on the held-out test set using the coefficient of determination (R²), mean absolute error (MAE), and root mean squared error (RMSE) for each response variable independently.

### Feature Importance Analysis

Feature importance was extracted from the Gini impurity-based importance scores of each individual gradient boosting sub-model. Importance values were analyzed both per-response-variable (to identify the dominant predictors for each regulatory layer) and averaged across all 16 response variables (to identify globally important features). Features were further grouped into six biological categories (structural/motif, spatial, baseline m6A, baseline poly(A), baseline isoform, baseline expression) and category-level importance was calculated as the sum of individual feature importances within each group, expressed as a percentage of total importance.

### Focused Validation of m6A Erosion Predictability Baseline Feature Computation

Baseline features were recomputed by pooling all Control reads per transcript across all Control samples. This pooling strategy provides stable estimates of intrinsic transcript properties by maximizing the number of observations per transcript. For each transcript, features were computed across six categories: m6A density (6 features: sites/kb, mean and median modification probability, number of sites, number of reads, detection rate), poly(A) tail statistics (15 features: distributional parameters including mean, median, standard deviation, coefficient of variation, percentiles, skewness, kurtosis, and proportions of short and long tails), motif composition (20 features: counts of 18 individual DRACH pentamers, motif diversity entropy, and DRACH percentage), spatial distribution (9 features: positional statistics including mean and median normalized position, position entropy, spatial clustering variance, three-prime enrichment, quartile counts, and mean distances from transcript start and end), and transcript structure (1 feature: transcript length). Treated samples (D7 and D14) were processed per-sample, and features were prefixed with “baseline_” for Control-derived predictors. The target variable was the change in m6A site density (Δsites/kb = treated sites/kb − baseline sites/kb).

### Gene-Grouped Cross-Validation

To prevent information leakage between transcripts of the same gene, 5-fold cross-validation was performed using GroupKFold (scikit-learn) with gene name as the grouping variable. This ensures that all transcripts from a given gene are assigned to the same fold, preventing the model from exploiting shared sequence features between transcripts of the same gene that appear in both training and test sets. Individual GradientBoostingRegressor models were trained within each fold (n_estimators=300, max_depth=4, subsample=0.8, random_state=42). Model performance was evaluated as the mean R² across all five folds, and out-of-fold predictions were collected for all transcripts.

### Feature Category Ablation

To determine the minimal feature set required for accurate prediction, models were trained using nested feature subsets: (A) baseline m6A density only (6 features), (B) m6A + poly(A) + structure (22 features), (C) B + motif composition (42 features), (D) B + spatial distribution (31 features), and (E) all features (51 features). Additionally, standalone models were trained using motif composition features only (20 features) and spatial distribution features only (9 features) to assess the independent predictive power of each category. All models were evaluated using gene-grouped 5-fold cross-validation as described above.

### Permutation Test

To formally test the null hypothesis that baseline features have no predictive relationship with m6A erosion, 100 permutation iterations were performed. In each iteration, the target variable (Δsites/kb) was randomly shuffled while preserving the feature matrix, and the full gene-grouped 5-fold cross-validation procedure was repeated. The empirical p-value was calculated as the proportion of permuted R² values that equaled or exceeded the observed R². The null distribution of permuted R² values was compared against the observed R² to assess statistical significance.

### Quantification of the Uniform Proportional Erosion Mechanism

The linear relationship between baseline m6A density and perturbation-induced m6A loss was quantified using ordinary least-squares regression (scikit-learn LinearRegression) and Pearson correlation. The regression slope provides a direct estimate of the proportional erosion rate.

Temporal dynamics were assessed by fitting separate regression models for D7 and D14 timepoints and comparing slopes.

### Response Pattern Clustering and Pathway Enrichment Correlation Analysis

To assess the degree of coordination among epitranscriptomic responses, Pearson correlation coefficients were computed pairwise among 15 response variables (the 16 described above, excluding expression log2FC to retain n=33,583 samples that would otherwise be lost to missing TPM data). The resulting 15x15 correlation matrix was visualized as a lower-triangular heatmap.

### Principal Component Analysis

The 15 response variables were standardized using z-score normalization (StandardScaler) and subjected to principal component analysis (PCA) to identify the major axes of variation in the multi-dimensional response space. A scree plot and cumulative variance explained curve were used to assess dimensionality. PCA loadings for the first three principal components were examined to characterize the biological interpretation of each axis.

### K-Means Clustering

Transcripts were clustered by their multi-dimensional response patterns using K-means clustering on the standardized 15-dimensional response vectors. The optimal number of clusters was determined by evaluating k=2 through k=10, using both the elbow method (within-cluster sum of squares) and silhouette analysis (mean silhouette coefficient). K-means was run with 10 random initializations (n_init=10) and a fixed random seed (random_state=42) for each value of k. Based on the convergence of both metrics, k=4 was selected as the optimal cluster number. Final cluster assignments were generated using these parameters, and cluster centers were back-transformed to the original response variable scale for biological interpretation.

### Signature Gene Identification

Signature genes for each cluster were identified using a distance-to-center approach. Prior to distance calculations, predicted genes (Gm-prefix followed by digits), RIKEN clones (Rik-suffix), and ribosomal protein genes (Rpl, Rps, Mrpl, Mrps prefixes) were removed from the gene set to prevent their over-representation from obscuring pathway-specific biology. Gene-level mean response vectors were computed by averaging response values across all transcripts and timepoints for each gene. For each gene, the Euclidean distance to each of the four cluster centers (back-transformed to the original 15-dimensional response variable scale) was computed, and the gene was assigned to its closest cluster center. The 550 genes closest to each cluster center were selected as signature genes for enrichment analysis.

### Pathway Enrichment Analysis

Gene lists for each cluster (after filtering) were submitted to the Enrichr API for pathway enrichment analysis. Five gene set libraries were queried: GO_Biological_Process_2023, GO_Molecular_Function_2023, GO_Cellular_Component_2023, KEGG_2021_Human, and Reactome_2022. Enrichment results included p-values, odds ratios, combined scores, and Benjamini-Hochberg adjusted p-values. A normalized pathway enrichment heatmap was generated to compare enrichment patterns across clusters, with p-values displayed on heatmap cells.

### Differential Gene Expression Analysis

Differential gene expression between Day 7 and Control conditions was assessed using DESeq2 [19]. Transcript-level quantification was performed using Salmon, and transcript abundances were summarized to gene-level counts using tximport with length-scaled TPM (countsFromAbundance = “lengthScaledTPM”) and a transcript-to-gene mapping derived from the GENCODE vM38 annotation. The count matrix comprised 4 Control and 3 Day 7 samples.

Low-count genes were pre-filtered by requiring a minimum of 10 counts in at least 3 samples. DESeq2 was run with default size factor estimation and dispersion estimation via DESeqDataSetFromTximport. Results were extracted using the Bor_D7 vs Control contrast, and log2 fold-change shrinkage was applied using the apeglm method for more accurate effect size estimates.

### Western Blot Analysis

DRG tissue was harvested from mice treated with bortezomib or vehicle at Day 5 and Day 9 post-treatment. Protein was extracted from the L4-6 DRGs in lysis buffer (50 mM Tris HCl, 1% Triton X-100, 150 mM NaCl, and 1 mM EDTA at pH 7.4) containing protease and phosphatase inhibitor mixtures with an ultrasonicator on ice, and cleared of cellular debris by centrifugation at 14,000 relative centrifugal force for 15 µmin at 4°C. Fifteen micrograms of protein per well were loaded and separated by standard 7.5% or 10% sodium dodecyl sulfate polyacrylamide gel electrophoresis. Proteins were transferred to Immobilon-P membranes (Millipore Sigma, Cat # IPVH00010) and then blocked with 5% dry milk for 3 h at room temperature. The blots were incubated with primary antibodies OxPhos Rodent WB Antibody Cocktail (1:1000, overnight at 4°C, Thermo Fisher, Cat # 45-8099) and beta-III-tubulin (1:50,000, 20 minutes RT, Promega, Cat # G7121) and detected the following day with donkey anti-rabbit or goat anti-mouse antibody conjugated to horseradish peroxidase (1:10,000, Jackson Immunoresearch, Cat # 711-036-152, Cat # 115-036-062). Signal was detected by enhanced chemiluminescence. Densitometric analyses were done using UN-SCAN-IT 7.1 software (Silk Scientific Corp.).

### Gene-Level Case Study Analyses Per-Site Data Extraction

For in-depth case studies, per-site m6A modification data were extracted for individual genes of interest (*Ldha, Sdhb, Uqcrc2*) from the complete feature matrix (ml_features_complete.csv). Each extracted dataset retained single-molecule resolution, preserving read-level modification probabilities, poly(A) tail lengths, positional information, motif identity, and sample/group assignments. Isoform identity was parsed from m6Anet transcript identifiers using GENCODE nomenclature.

### Isoform Structure Visualization

Exon-intron structures for expressed isoforms were reconstructed from the GENCODE vM38 GTF annotation by extracting all exon coordinates for each transcript. Isoform structures were visualized as gene diagrams with proportionally scaled exons and introns, with distinct color assignments for each isoform maintained consistently across all figure panels.

### Isoform Usage Analysis

Isoform-level expression proportions were calculated from nanopore read counts assigned to each isoform within each experimental condition. Read counts per isoform were normalized to proportions within each sample, and stacked bar charts were generated to visualize condition-dependent isoform switching.

### Positional m6A Hotspot Analysis

To identify positional patterns of m6A modification change, each gene body was divided into 100 equal bins along the normalized transcript coordinate (0 to 1). The density of m6A sites within each bin was calculated as the percentage of total sites falling within that bin for each experimental group. Differential density was computed as the bin-wise difference between treatment (D7 or D14) and Control, revealing positional hotspots of m6A gain or loss. The dominant 5-mer motif at each position was identified as the most frequently occurring kmer among sites within each bin.

### Motif and RBP Binding Analysis

Position-resolved motif analysis was performed by constructing gene-length × motif heatmaps showing the change in m6A modification density for each of the 18 DRACH 5-mer motifs at each positional bin. This analysis was conducted at both the gene level (aggregating across all isoforms) and the isoform level (separately for each expressed isoform). RNA-binding protein (RBP) motifs were obtained from the ATtRACT database [9], filtered to Homo sapiens and Mus musculus entries. For each DRACH motif, matching RBPs were identified by exact substring matching between the 5-mer motif and ATtRACT binding motif sequences (with U-to-T conversion). Position-resolved RBP heatmaps were generated by mapping RBP binding site changes across the transcript body, revealing spatially coordinated loss or gain of specific RBP recognition sites upon treatment.

### Poly(A) Tail Analysis

Single-molecule poly(A) tail length distributions were compared across experimental conditions using violin plots. Per-molecule poly(A) lengths were extracted from the integrated feature matrix for each gene, preserving individual read-level measurements. Statistical significance was assessed using the Kruskal-Wallis test (for multi-group comparisons) and Mann-Whitney U test (for pairwise comparisons), with effect sizes reported as the difference in group medians.

### Single-Cell Integration Analysis Single-Nuclei RNA Sequencing Data

Nuclei were extracted from DRGs of naïve male ICR mice using the Chromium Nuclei Isolation Kit (10x Genomics). Single-nuclei libraries were prepared using the Chromium Next GEM Single Cell 3’ Kit (10x Genomics) on the Chromium X instrument. Libraries for single-nucleus nanopore sequencing were prepared using the Ligation Sequencing DNA V14 Kit (Oxford Nanopore Technologies, SQK-LSK114) and sequenced on the PromethION P2 Integrated (Oxford Nanopore Technologies) using R10 flow cells (Oxford Nanopore Technologies, FLO-PRO114M). The wf-single-cell pipeline (Oxford Nanopore Technologies) was used for barcode demultiplexing, UMI deduplication, and generation of the gene–cell count matrix. The processed gene expression matrix was loaded from the 10x-format output (features.tsv.gz, barcodes.tsv.gz, matrix.mtx.gz).

### Cell Clustering and Annotation

Single-cell analysis was performed using Scanpy (v1.9). The standard preprocessing pipeline included: quality control metric calculation, total-count normalization to 10,000 counts per cell (normalize_total, target_sum=1e4), log1p transformation, selection of the top 2,000 highly variable genes, PCA using the ARPACK solver, construction of a k-nearest neighbor graph (n_neighbors=10, n_pcs=40), UMAP dimensionality reduction, and Leiden community detection clustering (resolution=0.5). Cell types were annotated based on the expression of established DRG marker genes [4]: nociceptors (Calca, Tac1, Trpv1, P2rx3, Mrgprd, Scn9a, Scn10a, Trpa1, Ntrk1), mechanoreceptors (Ntrk2, Ntrk3, Sst, Pvalb), Schwann cells (Mbp, Mpz, Plp1, Pmp22), satellite glia (Fabp7), and fibroblasts (Col1a1, Col1a2, Dcn). Clusters were assigned by applying marker score thresholds: clusters with >15% combined nociceptor marker expression as nociceptors, >10% combined mechanoreceptor markers as mechanoreceptors, >20% Schwann marker expression as Schwann cells, >10% Fabp7 as satellite glia, >5% fibroblast markers as fibroblasts, and clusters with >10% Pecam1 or Egfl7 >10% as endothelial cells.

### m6A Vulnerability Mapping

To project bulk epitranscriptomic vulnerability onto the single-cell landscape, genes assigned to the OXPHOS-enriched response cluster (Cluster 3) from the bulk K-means analysis were mapped to the single-cell gene expression matrix by gene symbol matching. Gene-level coverage was assessed as the percentage of bulk Cluster 3 genes detected in the single-cell matrix.

Two complementary vulnerability metrics were computed per cell. The compositional metric quantified the fraction of each cell’s total UMI counts attributable to Cluster 3 genes (cluster3_umis / total_umis). The magnitude-weighted burden metric incorporated the actual magnitude of m6A loss by computing, for each cell, the sum of UMI count × mean Δm6A (sites/kb) across all expressed Cluster 3 genes. This metric weights cells not only by how many vulnerable genes they express but also by how heavily each gene is affected by m6A erosion. The sign was inverted (multiplied by −1) so that higher values indicate greater m6A loss burden.

### Cell-Type Vulnerability Statistics

Per-cell vulnerability scores were compared across Leiden-defined cell types. Cells were grouped into sensory neurons (nociceptors and mechanoreceptors) versus non-neuronal populations (Schwann cells, satellite glia, fibroblasts, endothelial cells). Statistical comparison was performed using Welch’s t-test (for unequal variances) and the Mann-Whitney U test (non-parametric), with Cohen’s d calculated as the effect size measure. UMAP embeddings were colored by magnitude-weighted m6A loss burden to visualize the spatial distribution of vulnerability across cell populations.

### Software and Computational Environment

Basecalling was performed using Dorado 0.9.6. Alignment used minimap 2.24-r1122. m6A modification detection used m6Anet 2.1.0. Isoform analysis used FLAIR 3.0.0. Single-cell analysis used Scanpy. Differential expression analysis used DESeq2 with apeglm for log2 fold-change shrinkage. Machine learning was implemented in Python using scikit-learn (GradientBoostingRegressor, MultiOutputRegressor, StandardScaler, KMeans, PCA), with data manipulation in pandas and NumPy, and statistical testing via SciPy. Pathway enrichment was performed via the Enrichr web API. Visualization used matplotlib and seaborn. Reference genome: GRCm39 (mm39). Gene annotation: GENCODE vM38.

## RESULTS

### Baseline transcript features predict m6A loss with near-perfect accuracy but fail to predict other transcriptomic dynamics

Among the more than 170 known RNA modifications, N6-methyladenosine (m6A) is the most abundant internal modification of mammalian mRNA and plays critical roles in virtually every aspect of RNA metabolism[17]. m6A is deposited co-transcriptionally by the METTL3/METTL14 methyltransferase complex within a DRACH consensus motif (D = A/G/U, R = A/G, H = A/C/U) and is removed by the demethylases FTO and ALKBH5. m6A reader proteins, including the YTH domain family (YTHDF1/2/3, YTHDC1/2) and IGF2BP family (IGF2BP1/2/3), interpret the m6A code to direct transcript fate decisions including translation efficiency, mRNA decay, and localization[17]. Dysregulation of m6A has been implicated in cancer[38], neurological diseases, and immune disorders[39]. However, epitranscriptomic regulation of mRNA fate extends well beyond any single modification. Poly(A) tail length independently modulates mRNA stability and translational competence through interactions with poly(A)-binding proteins[1], while alternative isoform usage can reshape protein function by including or excluding functional domains[25].

Although each of these layers has been implicated in disease, whether intrinsic transcript features predetermine how any of them respond to perturbations has not been investigated.

To test whether drug-induced epitranscriptomic changes are deterministic consequences of intrinsic transcript architecture or arise from dynamic, context-dependent regulatory processes, we performed nanopore direct RNA sequencing on DRG harvested at Days 7 and 14 from mice treated with bortezomib for five consecutive days or vehicle controls and trained a multi-output gradient boosting model to simultaneously predict 16 molecular response variables spanning m6A remodeling, poly(A) tail dynamics, isoform switching, and expression change (Table S1) from 69 baseline transcript features (Table S2). Two design choices deliberately maximized biological heterogeneity to stress-test the model: (1) outbred ICR mice introduced genetic diversity[33] that would obscure any strain-specific epitranscriptomic wiring, and (2) response variables were pooled across both timepoints, forcing the model to discover architecture-response relationships that generalize across early (Day 7) and established (Day 14) treatment phases rather than fitting phase-specific patterns. Baseline features measured exclusively in Control samples spanned six categories: structural/motif properties (transcript length, 18 DRACH motif counts), spatial features (positional distribution of m6A sites), baseline m6A metrics (modification density and probability), baseline poly(A) statistics (tail length distribution parameters), baseline isoform complexity (diversity, productivity), and baseline expression level (Control TPM). Treatment-timepoint measurements were used strictly as response variables, never as predictors, ensuring that predictions reflect intrinsic vulnerability rather than confounded treatment effects.

The model revealed a striking asymmetry in predictability across regulatory layers (Figure 1A). The change in m6A site density from Control (Δsites/kb) was predicted with near-perfect predictive fit (R² = 0.949), indicating that 94.9% of the variance in perturbation-induced m6A loss is explained by baseline transcript features alone. This result establishes that m6A erosion is not a stochastic consequence of drug treatment but a deterministic outcome encoded in the transcript’s pre-existing architecture. In contrast, all eleven poly(A) tail response variables were poorly predicted (R² = 0.093–0.189), with predictions collapsing toward the population mean and failing to capture the substantial transcript-level variation in actual poly(A) responses. Isoform entropy change (R² = 0.351) and expression fold-change (R² = 0.377) showed weak-moderate predictability.

**Figure 1.**
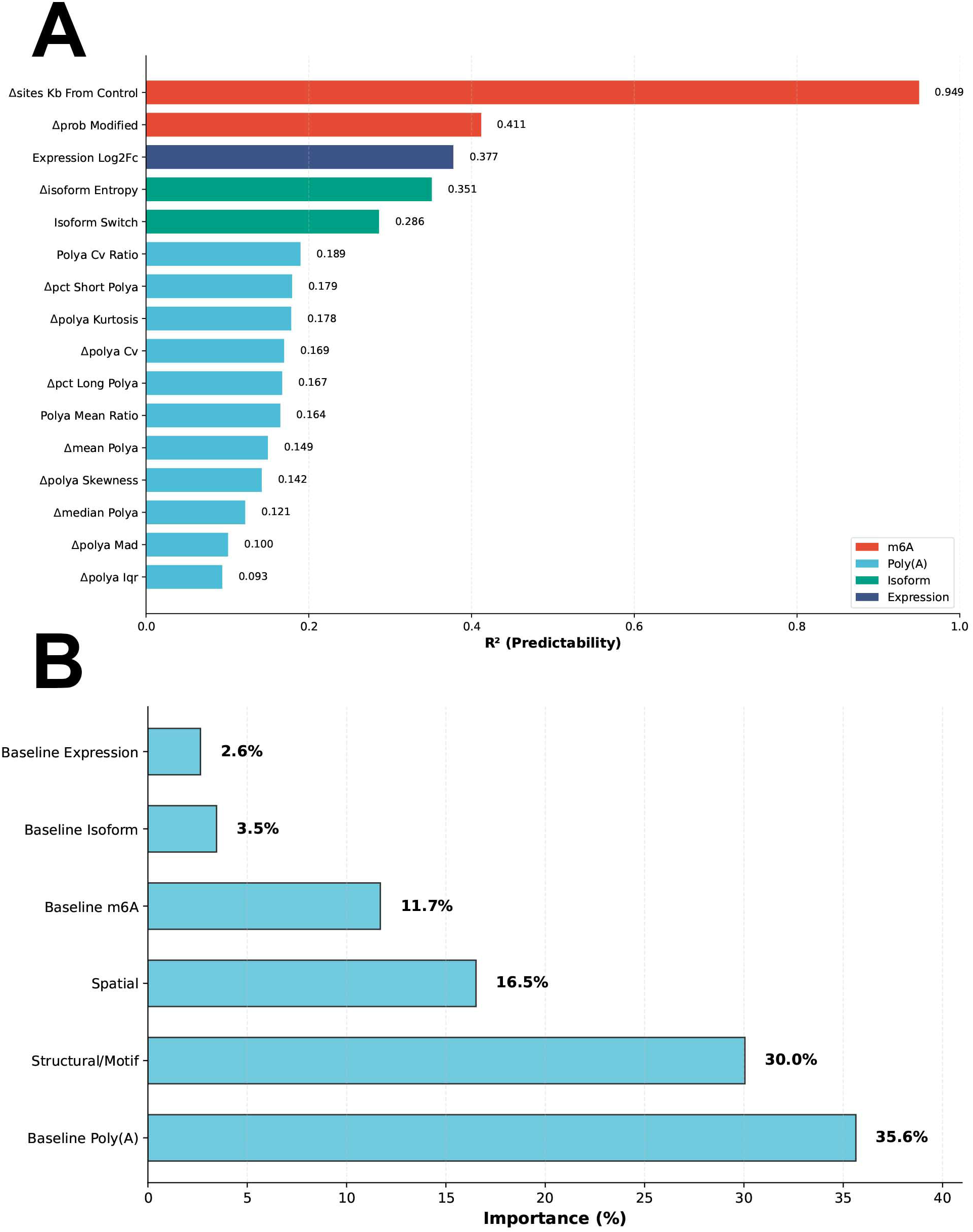
Machine learning prediction of perturbation-induced epitranscriptomic responses from baseline transcript features. (A) Model performance (R²) on the held-out test set for each of the 16 response variables, color-coded by regulatory layer: m6A (red), poly(A) (light blue), isoform (teal), and expression (dark blue). (B) Mean feature category importance averaged across all 16 response variable models. Sixty-nine baseline features were grouped into six categories.

Analysis of feature category importance averaged across all 16 response models revealed that baseline poly(A) features contributed the highest aggregate importance (35.6%), followed by structural/motif features (30.0%), spatial features (16.5%), baseline m6A features (11.7%), baseline isoform features (3.5%), and baseline expression (2.6%) (Figure 1B). The negligible contribution of expression, the sole metric accessible to conventional short-read transcriptomics, demonstrates that the dominant analytical paradigm in the field captures virtually none of the information that determines a transcript’s epitranscriptomic fate. The high importance of poly(A) features despite poor poly(A) model performance represents a notable dissociation. poly(A) baseline characteristics contribute to predicting responses in other layers, but the poly(A) response itself cannot be predicted from any combination of baseline features.

To examine these contrasts in detail, we compared the best-performing and worst-performing response models directly (Figure 2). The multi-output model was trained on 80% of transcript–timepoint observations and evaluated on a held-out 20% test set that the model never encountered during training. The predicted-versus-actual scatter plot for m6A site density loss on this held-out set (Figure 2A) confirmed that baseline features captured the full dynamic range of the response, with predictions tightly tracking actual values along the diagonal (R² = 0.949).

**Figure 2.**
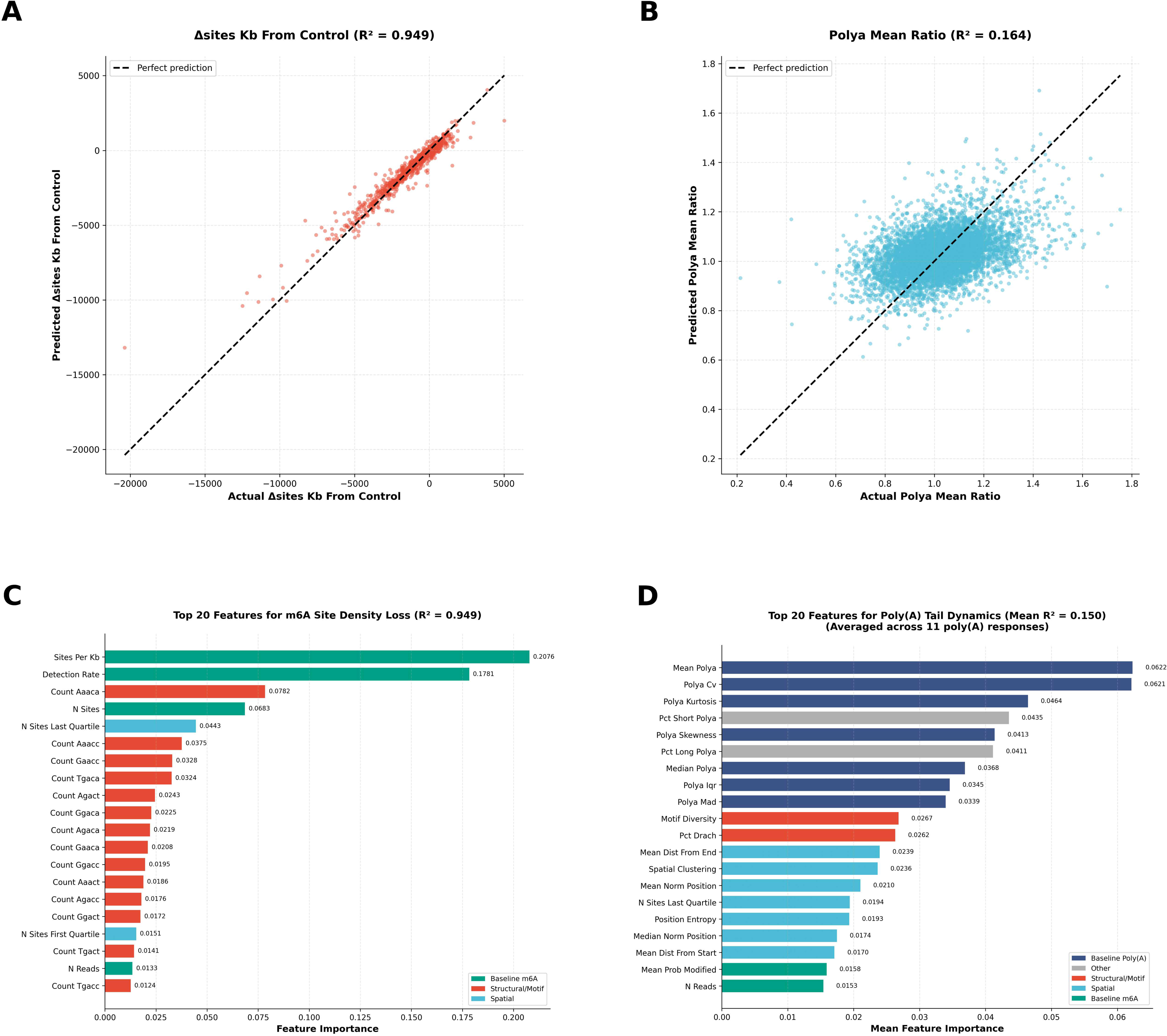
Prediction accuracy and feature importance for m6A and poly(A) models. (A) Predicted versus actual Δsites/kb on the held-out test set; dashed line = perfect prediction. (B) Predicted versus actual poly(A) mean ratio on the same test set. (C) Top 20 features by importance for the m6A site density loss model, color-coded by feature category. (D) Top 20 features by mean importance averaged across all 11 poly(A) response models, color-coded by feature category.

In stark contrast, the corresponding scatter plot for poly(A) mean ratio (Figure 2B) revealed predictions collapsing toward the population mean, failing to capture the substantial transcript-level variation in actual poly(A) responses (R² = 0.164). Predicted-versus-actual plots for all remaining response variables, including expression fold-change, m6A modification probability loss, all eleven poly(A) metrics, isoform entropy, and isoform switching, are shown in Figure S1. Feature importance analysis reinforced this asymmetry. The m6A site density loss model (Figure 2C) was dominated by baseline m6A site density (sites_per_kb, importance = 0.208) and detection rate (0.178), which together accounted for approximately 39% of total model importance, with fourteen of the top 20 features being specific DRACH pentamer motif counts (e.g., count_AAACA, importance = 0.078). The model thus predicts m6A loss primarily from the number and sequence composition of modifiable DRACH sites, transcript-intrinsic features defined by the RNA sequence itself, independent of whether those sites are actually methylated. In contrast, the poly(A) models (Figure 2D, mean R² = 0.150) showed importance distributed diffusely across baseline poly(A) statistics, structural features, and spatial features with no single dominant predictor, consistent with a response governed by factors external to intrinsic transcript architecture. Complete, feature importance profiles for expression change, m6A modification probability loss, and isoform switching models are provided in Figure S2.

### Focused validation confirms m6A erosion is encoded in transcript-intrinsic features

Given the striking predictability of m6A erosion revealed by the unbiased multi-output screen, we subjected this finding to rigorous focused validation. Three concerns warranted investigation: whether the high R² reflects genuine biological predictability rather than statistical overfitting, whether the model generalizes to genes not seen during training, and whether the relationship can be reduced to a simple mechanistic principle. To address these, we re-trained the m6A erosion model using gene-grouped 5-fold cross-validation, in which all transcripts from a given gene are assigned to the same fold, eliminating the possibility that sequence-similar transcripts from the same gene appear in both training and test sets. Baseline features were computed from pooled Control reads per transcript rather than per-sample averages, providing more stable estimates of intrinsic transcript properties.

Under this stricter validation regime, the m6A erosion model achieved R² = 0.983 ± 0.002 across five folds (Figure S3A), with performance stable across all folds (range: 0.980–0.985; Figure S3B). The improvement over the original held-out test set performance (R² = 0.949) reflects cleaner baseline estimation from pooled reads rather than reduced stringency, as gene-grouped cross-validation is a more conservative evaluation framework. Model residuals were centered near zero (mean = 0.817, std = 742.163 Δsites/kb) with no systematic bias (Figure S3C). The residual standard deviation of 742 Δsites/kb represents 1.2% of the observed range (0 to −60,000 Δsites/kb), confirming that prediction errors are small relative to the biological signal. Permutation testing (100 iterations with shuffled target labels) confirmed that the observed R² vastly exceeds the null distribution (p value approaching 0; Figure S3D), formally excluding the possibility that high-dimensional feature correlations produce spurious predictive accuracy.

Feature ablation revealed that predictive accuracy depends critically on baseline modification state. Sequence-intrinsic features alone, including pentamer counts and positional distributions, predicted m⁶A loss at R² = 0.113–0.164, while baseline m⁶A density alone achieved R² = 0.983 (Figure 3). This dissociation establishes that the predictive signal resides in the epitranscriptomic landscape, not nucleotide sequence, and is therefore invisible to conventional sequencing.

**Figure 3.**
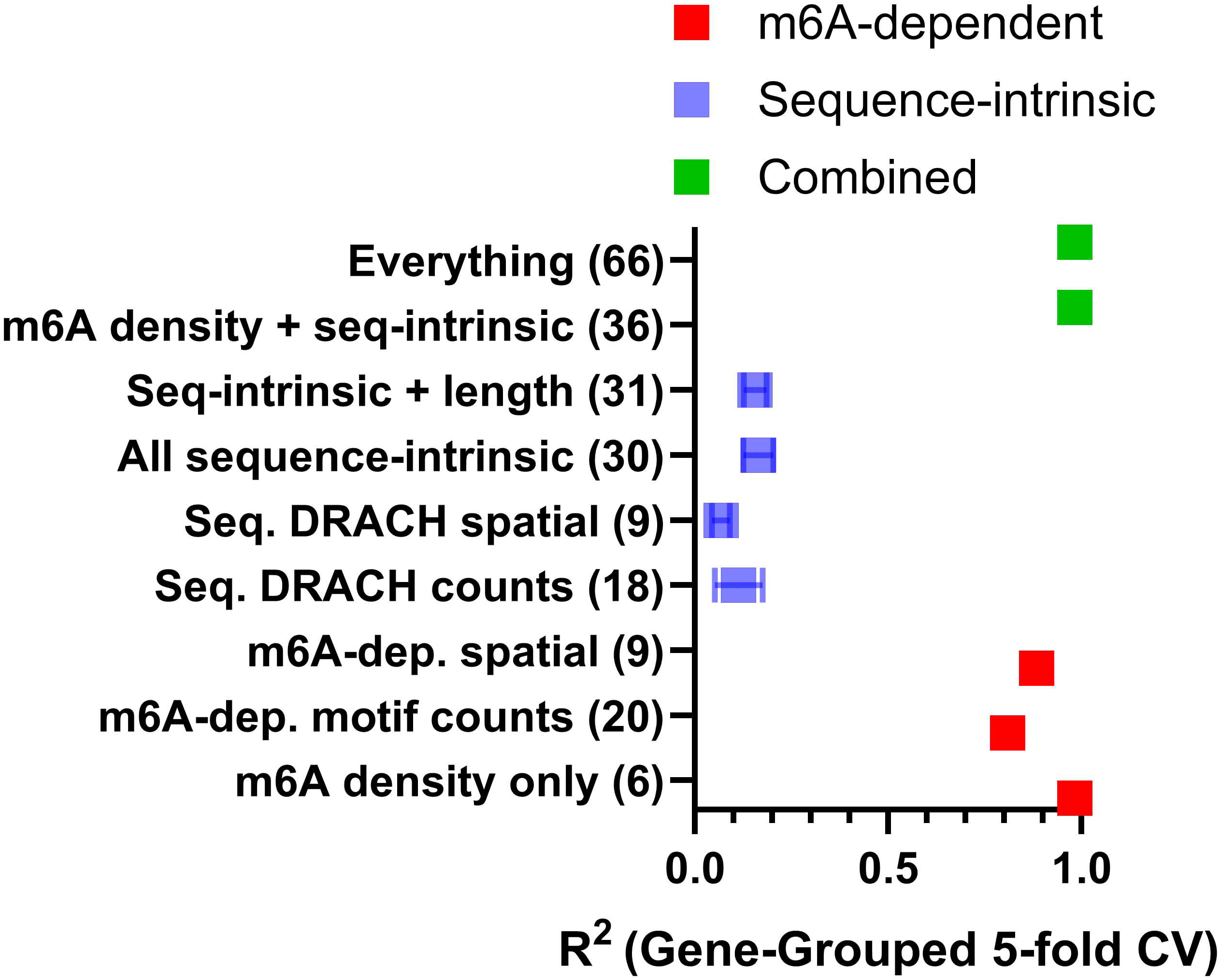
Feature ablation using gene-grouped 5-fold cross-validation. Numbers in parentheses indicate feature count per subset. m⁶A-dependent features (red) account for nearly all predictive performance, while sequence-intrinsic features (blue) contribute minimal signal. Combined models (green) shown for reference.

Baseline m6A density features (6 features) achieved the highest standalone performance (R² = 0.983; Figure 3), identical to the full 69-feature model (Figure S3E). Adding poly(A) tail statistics, transcript structure, and additional feature categories produced no further improvement (R² = 0.982 for all expanded models; Figure S3E). Gini importance analysis (Figure S3F) confirmed that m6A density features dominated when all categories were available (99.5% of total importance), consistent with m6A density being the most efficient, but not the only, encoding of the underlying architectural signal.

The relationship between baseline m6A density and perturbation-induced erosion followed a near-perfect linear relationship (Pearson r = −0.992, slope = −0.735; Figure S3G): for every 1,000 sites/kb of baseline m6A density, bortezomib treatment induces a loss of approximately 735 sites/kb. The constant proportional loss (∼73.5%) is consistent with a stoichiometric process in which a uniform fraction of m6A marks is removed regardless of local density. Temporal analysis revealed that the erosion fraction was greater at Day 7 (slope = −0.774, i.e., ∼77.4% loss) than at Day 14 (slope = −0.696, i.e., ∼69.6% loss; Figure S3H), indicating partial recovery of m6A marks during the later phase. Together, these results establish that m6A erosion follows a mechanistically simple proportional rule. The constant fractional loss means that a transcript’s absolute m6A erosion scales with its baseline density. The proportional kinetics thus describe how architecturally encoded vulnerability is realized while not being the source of the vulnerability itself.

### Epitranscriptomic regulatory layers operate independently

Beyond m6A methylation, poly(A) tail dynamics and alternative isoform usage represent additional layers of post-transcriptional regulation that can profoundly influence gene expression outcomes. Poly(A) tail length modulates mRNA stability and translational competence through interactions with poly(A)-binding proteins (PABPCs)[1], while alternative splicing and isoform switching can alter protein function by including or excluding functional domains[25]. Although mechanistic connections between individual regulatory layers have been described at specific loci, whether their responses to perturbation are coordinated across the transcriptome or operate as independent parallel mechanisms has not been systematically tested. Given the dramatic difference in predictability across regulatory layers, we next asked whether m6A, poly(A), and isoform responses are coordinated as part of an integrated epitranscriptomic program or operate as parallel, independent mechanisms. Pearson correlation analysis across 33,583 transcript–timepoint observations (Figure 4) revealed strong within-layer correlations, as expected: for example, delta_mean_polyA and delta_median_polyA were highly correlated (r = 0.83), and delta_polyA_cv and polyA_cv_ratio were nearly identical (r = 0.99). However, cross-layer correlations were uniformly near zero. m6A response variables showed correlations of |r| ≤ 0.06 with all poly(A) metrics, and isoform responses were essentially uncorrelated with both m6A and poly(A) changes (|r| ≤ 0.04).

**Figure 4.**
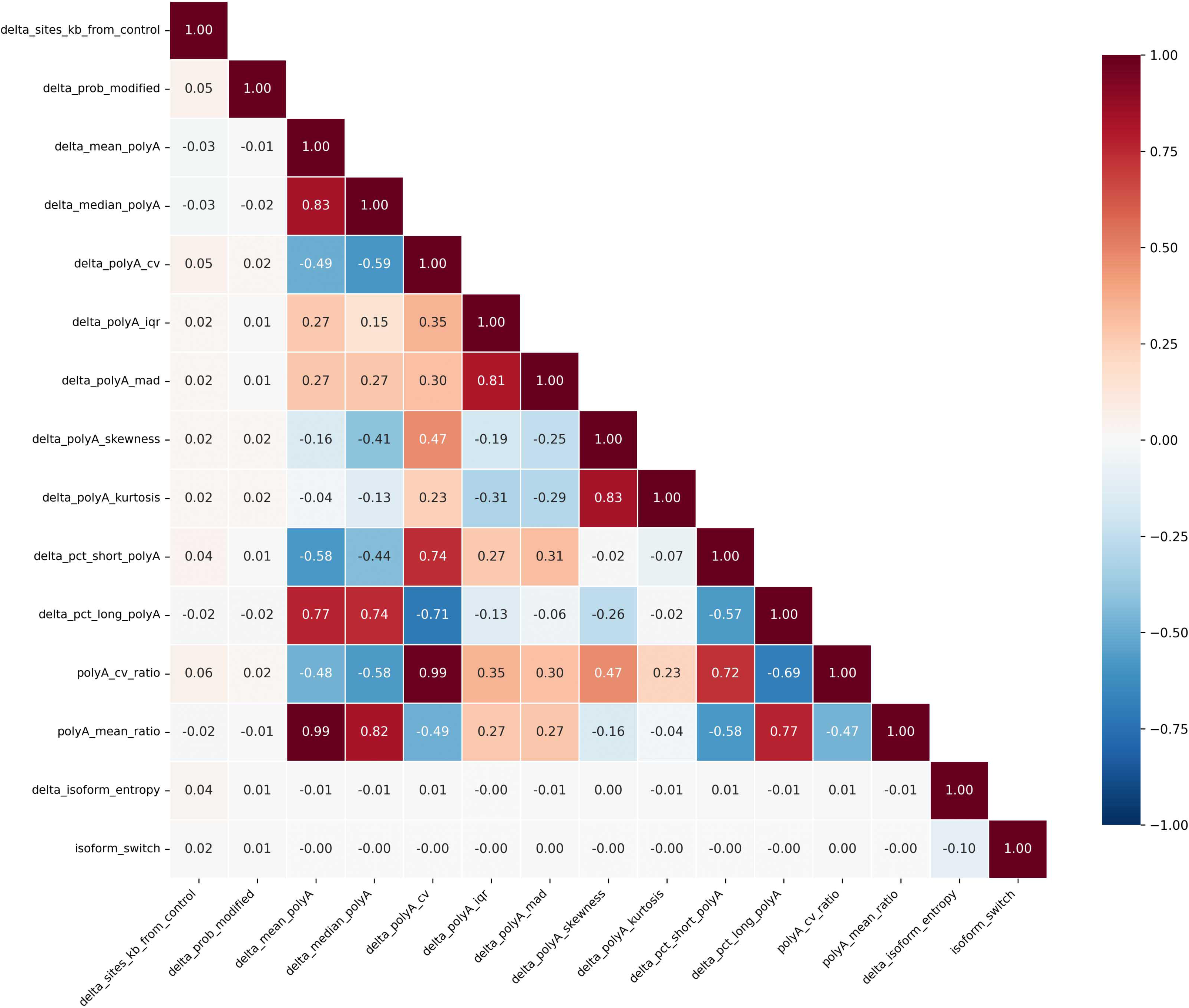
Pearson correlation matrix of 15 perturbation-induced response variables (expression excluded). Color scale: −1 (blue) to +1 (red).

Within this dataset, the near-complete independence of the three measured layers has two implications. First, no single upstream regulator appears to coordinately adjust m6A, poly(A), and isoform responses in this system, at least not through mechanisms detectable with the features captured here. Second, m6A and poly(A) responses appear to follow distinct regulatory logic: m6A erosion scales deterministically with baseline modification density, while poly(A) dynamics are poorly predicted by any measured transcript-intrinsic feature, suggesting dependence on trans-acting factors, signaling states, or regulatory inputs not captured by the current platform. Whether additional RNA modifications or transcript features not measured here mediate cross-layer coordination, or whether perturbation itself disrupts coordination that exists under homeostatic conditions, remains to be determined.

### Unsupervised clustering independently confirms m6A erosion as the dominant response signature

The gradient boosting model identified baseline m6A site density as the dominant predictor of epitranscriptomic response, but as an ensemble method its internal representation of response patterns is not directly interpretable. To independently test whether m6A erosion emerges as the predominant response signature through an unsupervised and fully interpretable framework, we performed K-means clustering on the standardized 15-dimensional response vectors (15 epitranscriptomic response variables, excluding expression to retain n = 33,583 observations). The optimal cluster number (k = 4) was determined by the convergence of the elbow method and silhouette analysis. The four clusters defined biologically interpretable response patterns (Table 1), with the largest cluster (Cluster 3; 43.5% of transcripts) defined by pronounced m6A site density loss (−487 sites/kb) with minimal poly(A) or isoform perturbation, confirming that m6A erosion is the most widespread epitranscriptomic consequence of this perturbation in DRGs.

### The m6A erosion cluster enriches for oxidative phosphorylation and proteasome pathway genes

To determine whether the empirically-defined response clusters correspond to coherent biological programs, we performed pathway enrichment analysis on the signature genes of each cluster (Figure 5). Cluster 3 (m6A erosion) showed overwhelming and exclusive enrichment for oxidative phosphorylation (p = 1.0 × 10⁻¹⁷) and proteasome-mediated proteolysis (p = 2.8 × 10⁻¹⁰). No other cluster showed significant enrichment for the OXPHOS pathway. The isoform switching cluster (Cluster 2) was enriched for mRNA splicing (p = 0.001) and modestly enriched for proteasome-mediated proteolysis (p = 0.03), while Cluster 0 (poly(A) elongation) showed modest enrichment for mitochondrion organization (p = 0.01).

**Figure 5.**
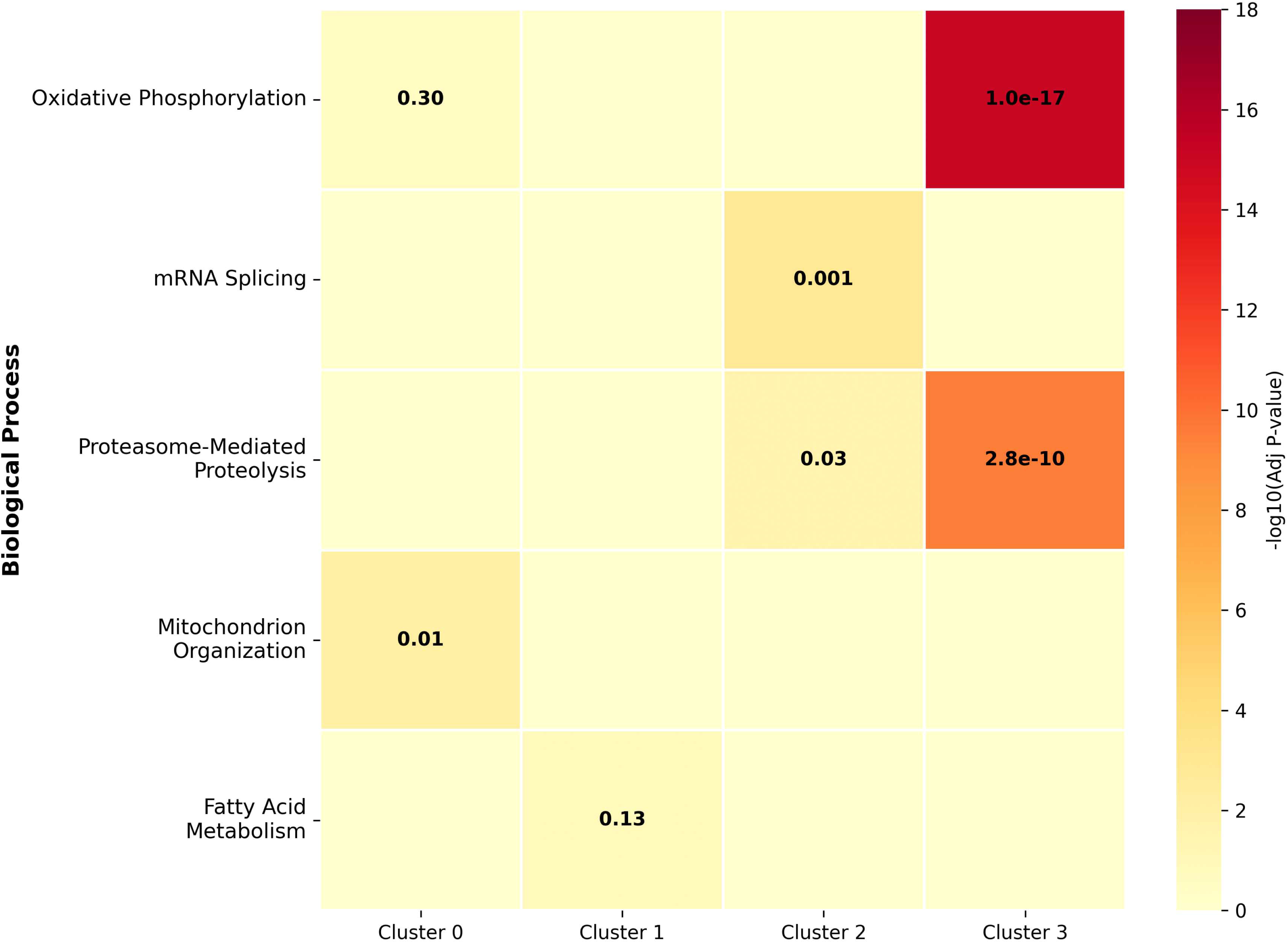
Cluster-specific pathway enrichment heatmap showing −log₁₀(adjusted p-value) for five biological processes across the four empirically defined response clusters.

This result is striking because it was obtained through a purely data-driven, unsupervised workflow that had no prior knowledge of bortezomib’s mechanism of action. The supervised model (Figure 2) demonstrated that intrinsic transcript architecture predetermines m6A vulnerability, and now the unsupervised clustering reveals that these architecturally determined responses converge on specific biological programs. These two independent computational approaches reach the same conclusion from different directions. One predicts individual transcript fates from baseline architectural features, while the other discovers emergent response patterns in an entirely unbiased manner. Yet both independently recapitulate the established mechanistic pathway of bortezomib-induced metabolic reprogramming, from proteasome inhibition and HIF1α stabilization[21] to the shift from oxidative phosphorylation toward aerobic glycolysis[20]. Purely computational analysis of intrinsic transcript architecture thus arrives at the same conclusions that required years of targeted cell and molecular biology experiments, validating that the architectural features encoded in each transcript carry biologically meaningful regulatory information.

Within the Cluster 3 signature gene set, key metabolic enzymes previously implicated in the bortezomib–HIF1α axis were prominently represented. LDHA, which catalyzes the terminal step of aerobic glycolysis, was assigned to this cluster. Numerous electron transport chain subunits from all five OXPHOS complexes—including NADH dehydrogenase subunits (Complex I), succinate dehydrogenase subunits such as *Sdhb* (Complex II), ubiquinol-cytochrome *c* reductase subunits such as *Uqcrc2* (Complex III), cytochrome *c* oxidase subunits (Complex IV), and ATP synthase subunits (Complex V)—underwent coordinated m6A erosion (Table S3). This pattern suggests that m6A erosion may facilitate the transcriptional and translational remodeling required for the metabolic switch, potentially by altering mRNA stability, ribosome recruitment, or interactions with m6A reader proteins at these metabolic transcripts.

### Differential gene expression analysis reveals minimal transcriptional changes at metabolic pathway genes

To directly test whether the conventional expression-centric analytical paradigm would detect the epitranscriptomic remodeling described above, we performed differential gene expression analysis on Day 7 versus Control samples. Strikingly, only 11 genes reached statistical significance after multiple testing correction: *Cfap100, Tcf4, Gm4117, Myt1, Atp11a, Pcdh1, Ybx1, Ybx3, Qki, Ank3,* and *Arf6*. Notably, none of the oxidative phosphorylation genes, glycolytic enzymes, or proteasome pathway components that populate the m6A erosion cluster showed significant changes in mRNA abundance.

This finding underscores the central insight of architectural determinism. A conventional SBS transcriptomic study of this system would conclude that bortezomib has minimal transcriptional impact on metabolic genes, entirely missing the extensive epitranscriptomic remodeling documented above. Yet functional and protein-level alterations in the bortezomib response are well documented. The near-absence of transcriptional changes at metabolic pathway genes, despite robust m6A remodeling at the same genes, establishes that the biological response is driven by post-transcriptional regulation encoded in transcript architecture rather than by classical transcriptional control. The machine learning model’s assignment of just 2.6% predictive importance to expression (Figure 1B) is thus not merely a computational observation but a biological reality. Hence, mRNA abundance is the wrong metric for understanding this response.

Among the 11 differentially expressed genes, *Ybx1* and *Ybx3* are noteworthy as they encode Y-box binding proteins that function as RNA-binding proteins involved in m6A- dependent mRNA stability[24; 35] and translational regulation[28], respectively. *Qki*, an RNA-binding protein of the STAR family, regulates mRNA splicing and stability[13; 27]. The differential expression of these RNA-binding proteins raises the possibility that the perturbation indirectly reshapes the post-transcriptional regulatory landscape through a small number of trans-acting factors, while the broad epitranscriptomic remodeling on metabolic genes operates through the m6A-dependent, architecture-encoded mechanism described above.

### *Ldha* epitranscriptomic remodeling reveals position-specific m6A loss at PABP recognition sites

To characterize the epitranscriptomic response at single-gene resolution, we performed detailed multi-layer analysis of *Ldha*, a key effector of the HIF1α-driven glycolytic switch and a Cluster 3 (m6A erosion) member. FLAIR isoform analysis identified five expressed *Ldha* isoforms in DRGs: *Ldha*-201, *Ldha*-210, *Ldha*-209, *Ldha*-214, and *Ldha*-202 (Figure 6A). The five *Ldha* isoforms varied in exon structure, with *Ldha*-201 and *Ldha*-210 differing in their 5′ exon architecture, *Ldha*-214 containing a distinct third exon, and Ldha-209 utilizing 7 exons with an alternative 3′ terminal exon compared to 8 in the other isoforms. Remarkably, isoform usage proportions were near-identical across all conditions (Figure 6B), with each of the four major isoforms maintaining 22.9–23.9% usage and Ldha-202 at 5.5–5.8% (maximum variation < 1 percentage point for any isoform). This stability rules out isoform switching as a mechanism for bortezomib-induced *Ldha* regulation.

**Figure 6.**
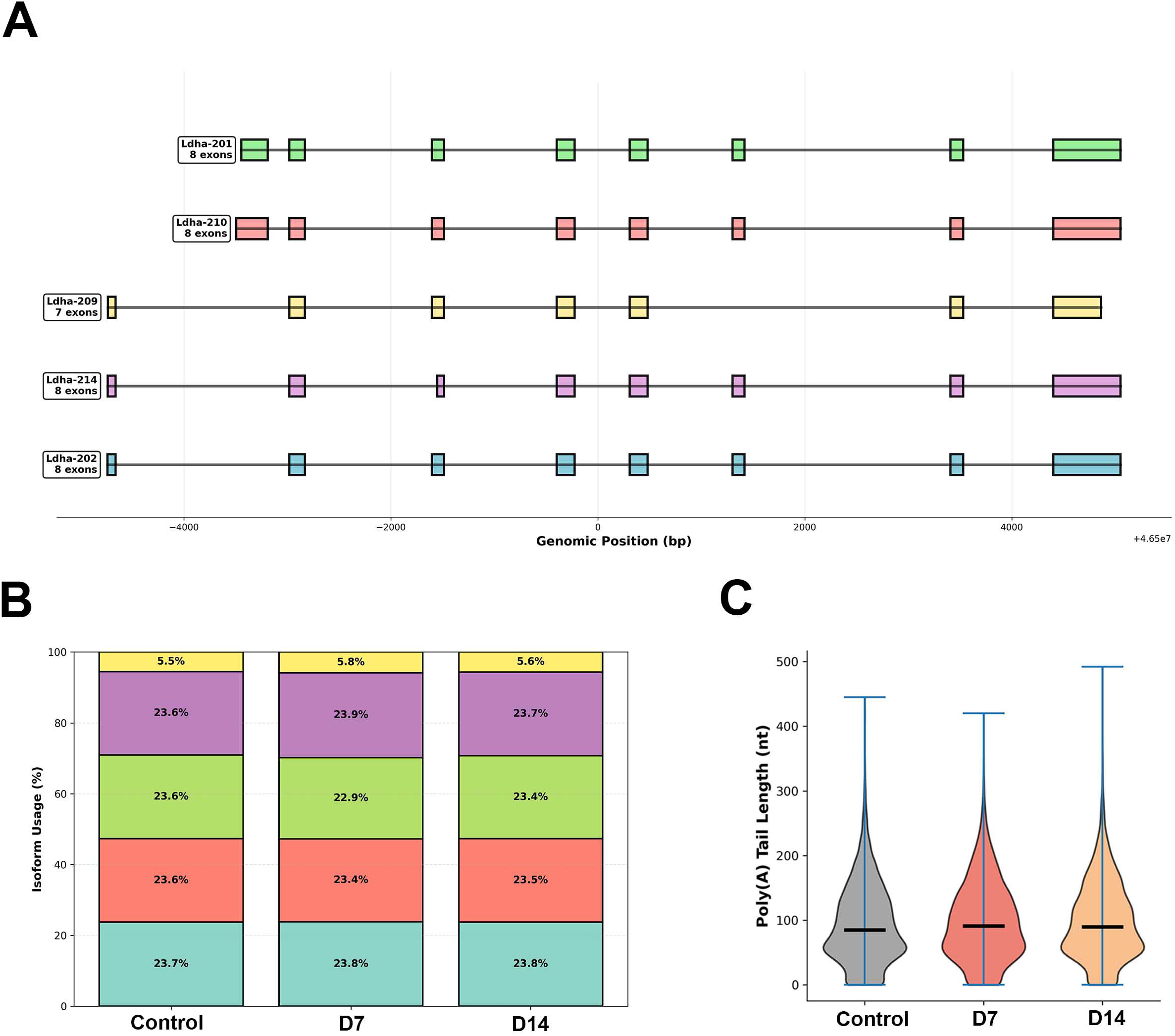
Multi-layer epitranscriptomic analysis of *Ldha*. (A) Exon-intron structures of the five *Ldha* isoforms expressed in mouse DRGs. Exons: filled rectangles; introns: connecting lines. (B) Isoform usage proportions across Control, D7, and D14 conditions. (C) Single-molecule poly(A) tail length distributions across conditions.

Single-molecule poly(A) tail analysis revealed a modest but statistically significant elongation of *Ldha* transcripts upon bortezomib treatment (Figure 6C). Median poly(A) length increased from 85 nt (Control, n = 9,438 reads) to 91 nt (D7, n = 4,189 reads) and 90 nt (D14, n = 6,179 reads). The Kruskal-Wallis test confirmed a significant global difference (H = 31.24, p = 1.65 × 10⁻⁷), with pairwise tests showing significance for Control vs. D7 (p_adj = 5.87 × 10⁻⁵) and Control vs. D14 (p_adj = 1.88 × 10⁻⁶) but not D7 vs. D14 (p_adj = 0.83). The ∼6 nt median shift in the poly(A) tail length was statistically significant, but the functional consequences of changes at this magnitude remain unclear.

Position-resolved motif analysis across the *Ldha* transcript body (Figure 7A), averaging across all five expressed isoforms, revealed an asymmetric m6A remodeling pattern that was motif-specific and temporally conserved. In the 5′ region, GAACA showed the most prominent gains at positions ∼10–20%, while several other motifs including TGACA and GAACC exhibited moderate gains through the 30–50% range. Past the 70% mark, m6A density declined broadly across nearly all 12 DRACH motifs, producing a striking 3′ loss wall that was not restricted to a single sequence context. AAACC and AGACC motifs at the extreme 3′ end (positions 90–100%) showed the deepest losses, and AAACA exhibited pronounced loss at 70–80%, while the overall pattern was conserved between D7 and D14.

**Figure 7.**
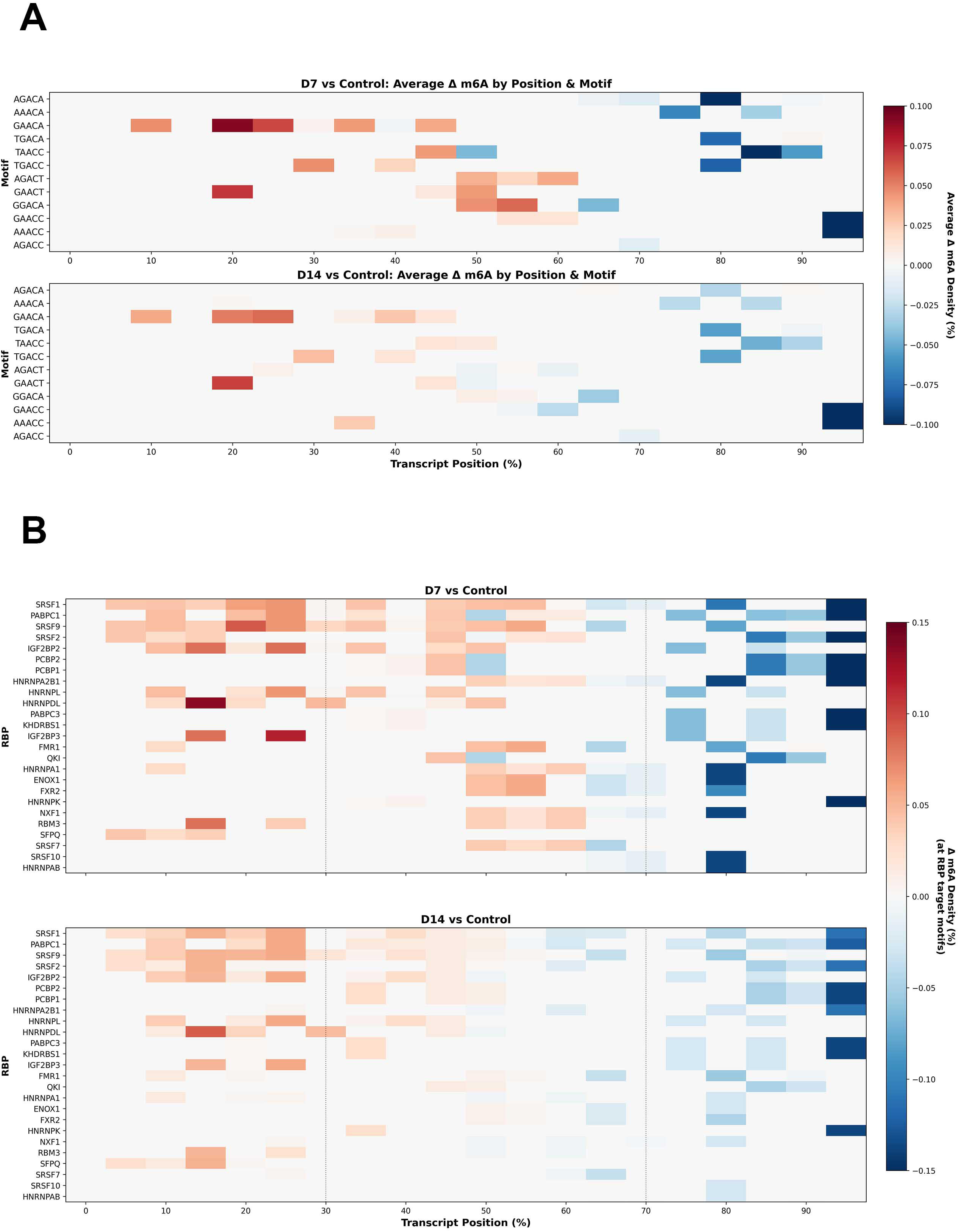
Position-resolved epitranscriptomic analysis of *Ldha* averaged across all five expressed isoforms. (A) Δm6A density (%) by DRACH motif identity (y-axis) and normalized transcript position (x-axis) for D7 vs Control (upper) and D14 vs Control (lower). Red: m6A gain; blue: loss. (B) Δm6A density at potential RBP target motifs across the same positional coordinates.

Mapping these positional changes onto RNA-binding protein target motifs (Figure 7B) revealed a concordant landscape across approximately 25 RBPs. The 3′ m6A loss affected recognition sites for a broad set of regulators, including PABPC1, PABPC3, HNRNPK, KHDRBS1, SRSF10, and HNRNPAB, while the 5′ region showed focal m6A gains at HNRNPDL sites (∼10–15%) and IGF2BP3 sites (∼30–40%). SRSF7 was notable for exhibiting m6A gain at positions ∼80–90%, running counter to the prevailing 3′ loss pattern. The 3′ loss intensified from D7 to D14 across the majority of RBPs. Among these affected sites, the PABPC1 and PABPC3 losses are of particular functional interest because AAACC is a recognition motif for poly(A)-binding proteins, key components of the cap-dependent translation initiation machinery[5; 29]. Reduced m6A at PABP binding sites in the 3′ UTR can potentially enhance PABP accessibility and thereby promote translation through the eIF4F–PABP closed-loop complex. This predicted translational enhancement is consistent with our previously published observations that bortezomib treatment increases LDHA protein levels in DRGs and elevates extracellular acidification rate (ECAR)[20], a direct measure of lactate extrusion. The m6A remodeling pattern at PABP recognition sites thus provides a potential post-transcriptional mechanism linking epitranscriptomic change to the increased glycolytic output observed in BIPN.

Isoform-resolved motif heatmaps (Figures S4A–E) showed that m6A losses in the 3′ region (past ∼70% of transcript length) were the most consistent feature of *Ldha* remodeling, present in *Ldha*-201 and -202 at both treatment timepoints. *Ldha*-210 showed strong 3′ losses at D7, though the pattern shifted toward the 5′ end by D14, while *Ldha*-214 displayed 3′ losses alongside more distributed gains in the first third of the transcript. *Ldha*-209 showed minimal remodeling at either timepoint. Modest 5′ m6A gains were observed across isoforms but were less consistent in position and magnitude. RBP analysis (Figures S5A–E), which mapped these m6A changes onto predicted RNA-binding protein recognition sites, produced consistent spatial profiles, identifying the specific RBPs likely affected by the positional remodeling at each isoform.

### m6A remodeling and post-transcriptional suppression of OXPHOS components

The observation that *Ldha* undergoes m6A loss at 3′ AAACC motifs raised the question of whether OXPHOS genes that are functionally suppressed during the metabolic switch might exhibit distinct epitranscriptomic remodeling. To explore this, we performed isoform-resolved analysis of *Sdhb*, which encodes the iron-sulfur subunit of mitochondrial Complex II, and *Uqcrc2*, which encodes Core protein 2 of mitochondrial Complex III.

In contrast to the multi-isoform architecture of *Ldha*, both *Sdhb* and *Uqcrc2* were expressed as single dominant isoforms in DRGs (*Sdhb*-201 and *Uqcrc2*-201, respectively), enabling unambiguous attribution of m6A changes to specific transcript positions. Position-resolved motif analysis revealed complex, position-dependent m6A remodeling at both genes. At *Sdhb* (Figure 8A), scattered m6A changes were observed across the transcript at D7, with a mixed pattern that included both gains and losses at various motifs and positions. By D14, the *Sdhb* landscape was dominated by m6A loss. RBP target analysis at *Sdhb* (Figure 8B) showed heterogeneous changes at PABP recognition sites, with evidence of increased m6A density at some positions but not a consistent transcript-wide pattern. At *Uqcrc2* (Figure 9A), D7 analysis revealed m6A gains at multiple motifs in the 3′ region of the transcript, including at AAACC and related PABP-associated sequences beyond the 90% transcript position. This 3′ gain pattern is partially consistent with a bidirectional model in which OXPHOS transcripts acquire m6A at sites where glycolytic transcripts like *Ldha* lose it, though the pattern was not uniformly present across both genes or timepoints. *Uqcrc2* RBP target analysis (Figure 9B) similarly revealed heterogeneous changes rather than the uniform gain pattern that a simple bidirectional mechanism would predict.

**Figure 8.**
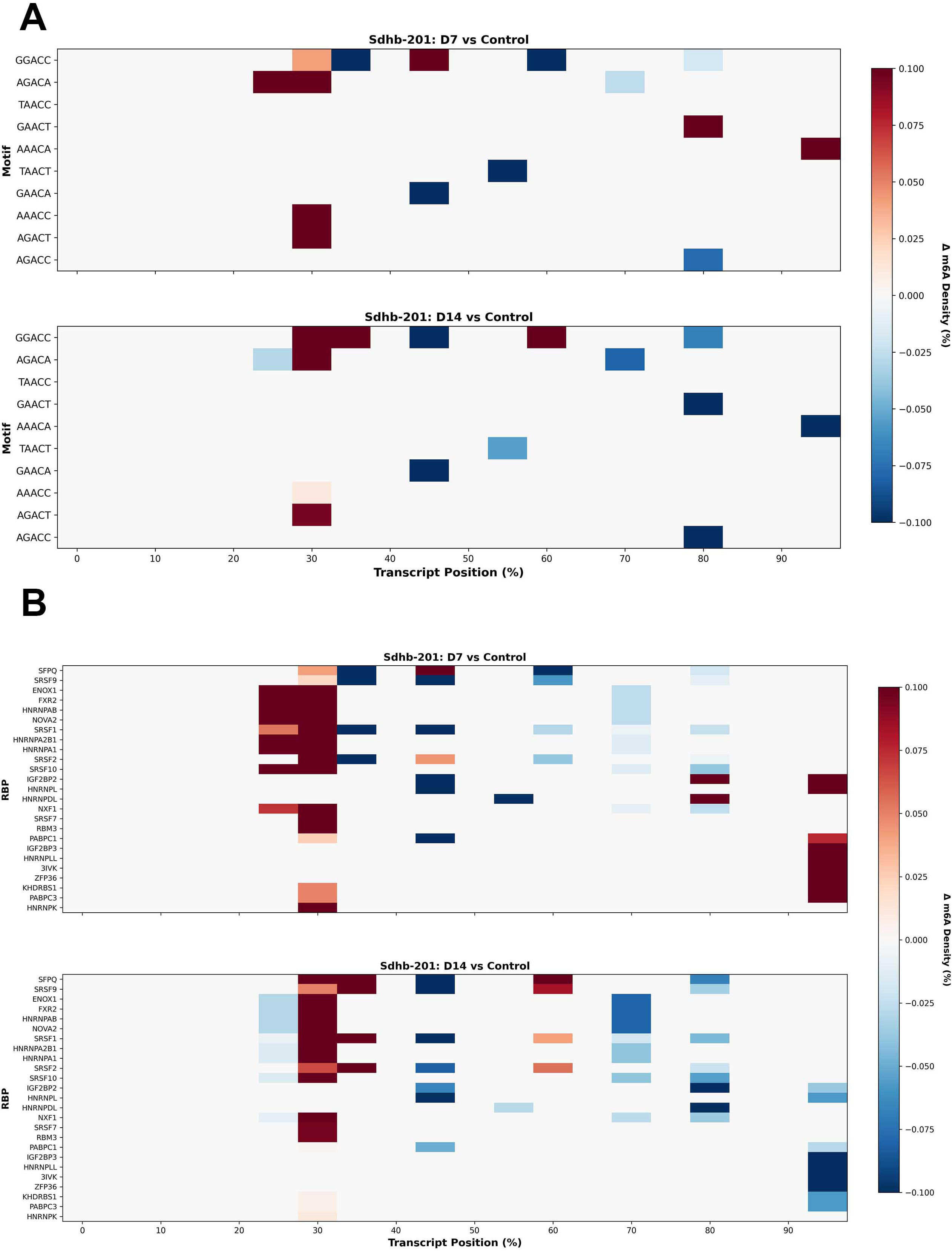
Position-resolved epitranscriptomic analysis of *Sdhb*-201. (A) Δm6A density (%) by DRACH motif identity and normalized transcript position for D7 vs Control (upper) and D14 vs Control (lower). Red: m6A gain; blue: loss. (B) Δm6A density at potential RBP target motifs across the same coordinates.

**Figure 9.**
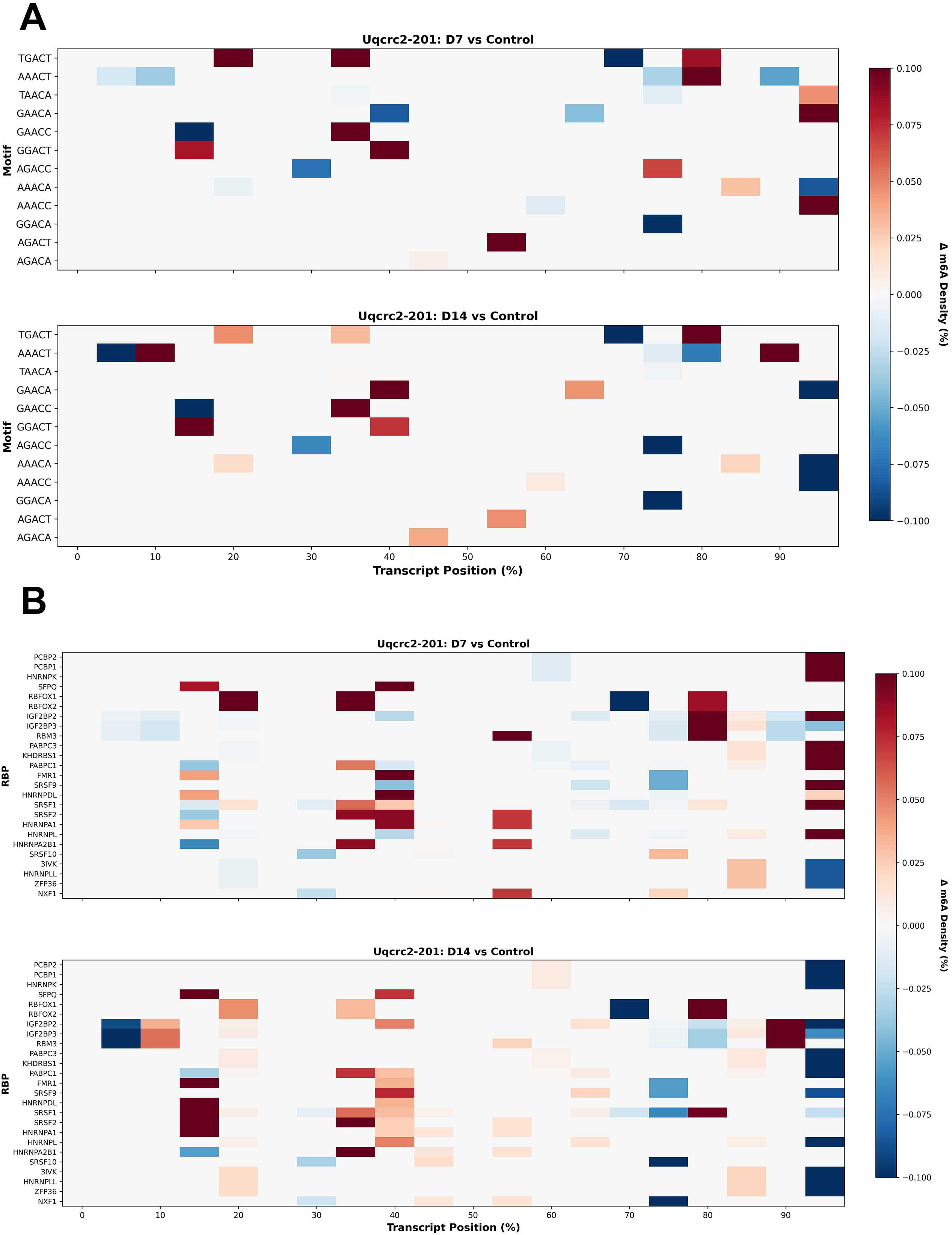
Position-resolved epitranscriptomic analysis of *Uqcrc2*-201. (A) Δm6A density (%) by DRACH motif identity and normalized transcript position for D7 vs Control (upper) and D14 vs Control (lower). (B) Δm6A density at potential RBP target motifs across the same coordinates.

To determine whether m6A remodeling at OXPHOS genes was associated with functional protein-level changes, we performed western blot analysis of DRG tissue. Western blot tissue was harvested at Day 5 and Day 9, timepoints that bracket the Day 7 sequencing harvest, allowing us to assess whether protein-level changes were already established before the epitranscriptomic measurement and whether they persisted beyond it. At Day 5, bortezomib treatment produced significant reductions in both UQCRC2 (p = 0.0028) and SDHB (p = 0.0020) protein levels relative to vehicle-treated controls, normalized to βIII-tubulin (Figure 10A-B).

**Figure 10.**
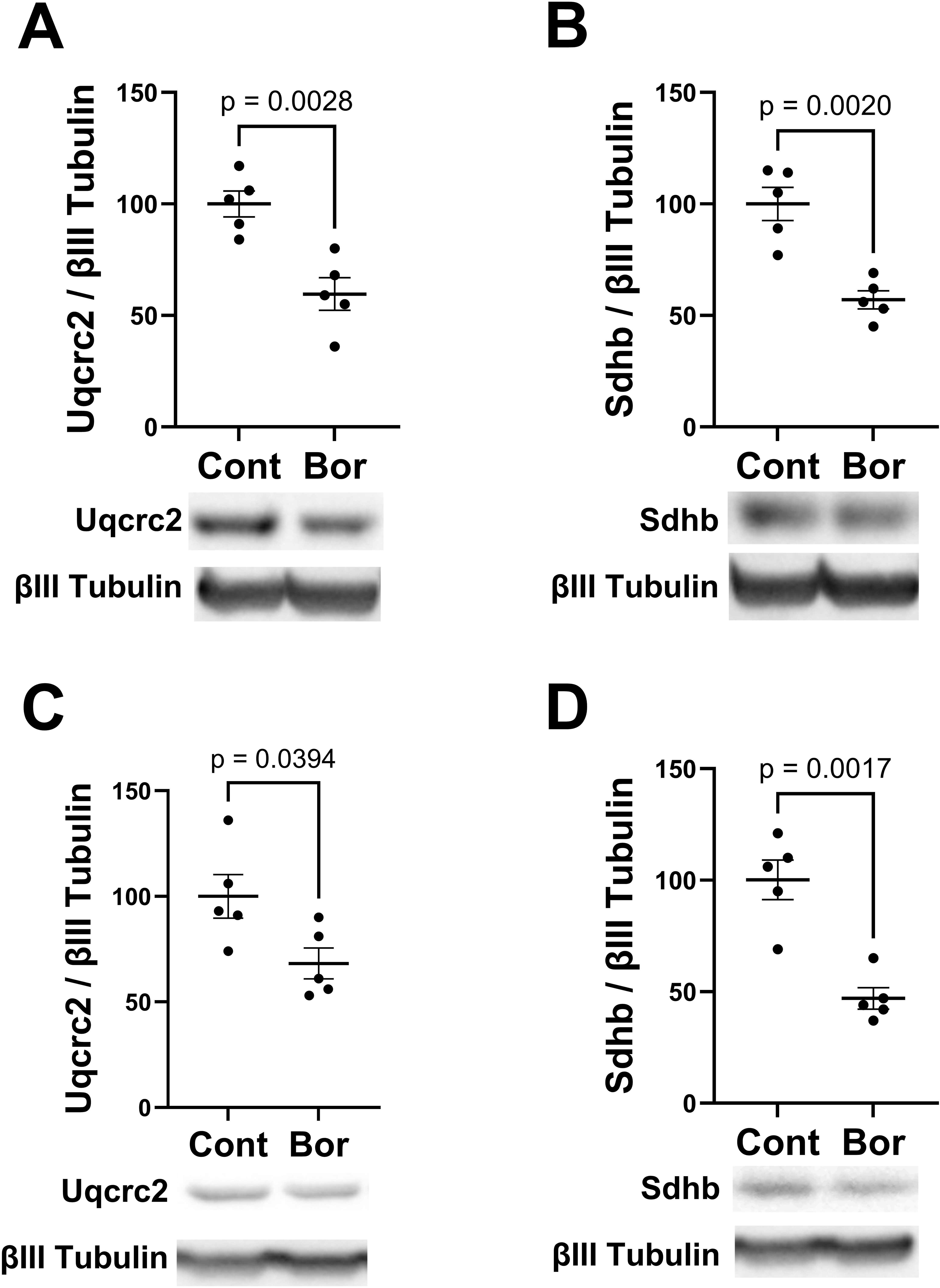
Western blot validation of OXPHOS protein suppression. Densitometric quantification and representative blots of UQCRC2 and SDHB normalized to βIII-tubulin. (A) UQCRC2 at Day 5. (B) SDHB at Day 5. (C) UQCRC2 at Day 9. (D) SDHB at Day 9. Data: mean ± SEM. Unpaired t-test.

These reductions persisted at Day 9 for both UQCRC2 (p = 0.0394) and SDHB (p = 0.0017; Figure 10C-D). The presence of protein suppression at both flanking timepoints indicates that the changes observed are not transient fluctuations but reflect a sustained shift in OXPHOS complex protein availability. Critically, neither *Sdhb* nor *Uqcrc2* showed significant changes in mRNA abundance by differential expression analysis, indicating that the protein-level suppression occurs through post-transcriptional regulation rather than transcriptional downregulation.

The decreased protein levels of Complex II and Complex III components are functionally concordant with the reduced oxygen consumption rates previously reported in DRGs from mice with BIPN [20] and independently corroborate the OXPHOS pathway enrichment rediscovered by unsupervised clustering of the epitranscriptomic data (Figure 5). The 3′ m6A gains at *Uqcrc2* are consistent with a model in which altered PABP accessibility contributes to translational suppression, but the heterogeneous remodeling landscapes across both genes and timepoints suggest that this might represent one component of a broader post-transcriptional program. As with *Ldha*, direct validation of the PABP mechanism remains to be established, and additional regulatory layers, including m6A reader protein recruitment and translational repression, likely contribute to the observed protein reductions.

### Sensory neurons carry disproportionately high m6A loss burden

The bulk analyses described above establish that OXPHOS genes undergo coordinated m6A erosion, but cannot distinguish whether this response occurs uniformly across DRG cell types or is concentrated in specific cell populations. To address this question, we integrated our bulk epitranscriptomic data with single-nuclei nanopore RNA sequencing of naive mouse DRGs.

Leiden clustering of 521 cells identified seven cell types based on established marker gene expression[4]: nociceptors, mechanoreceptors, Schwann cells, satellite glia, endothelial cells, fibroblasts, and a mixed population (Figure 11A, Table S4).

**Figure 11.**
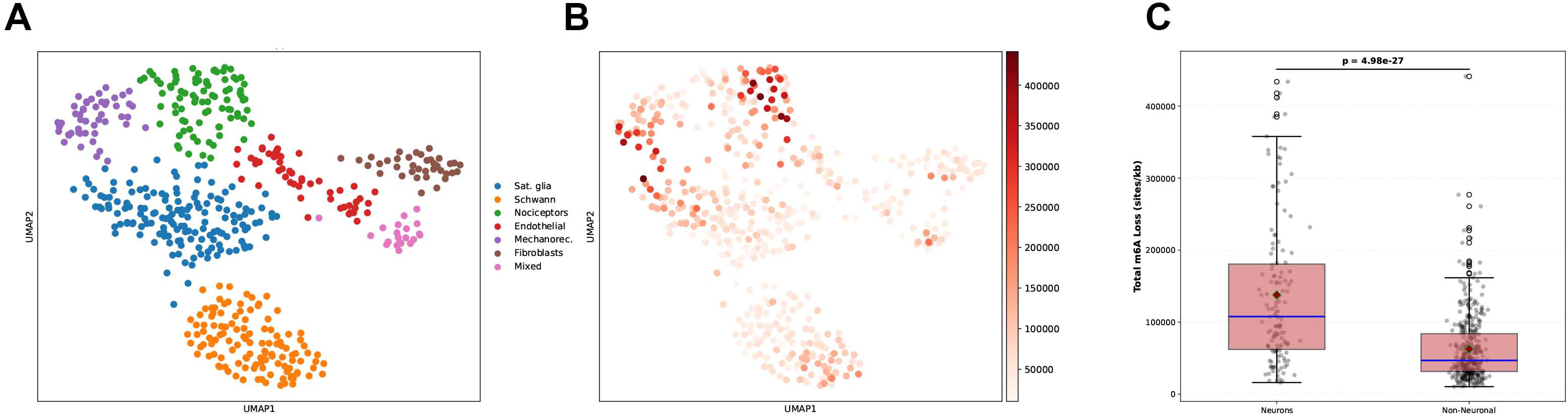
Single-cell m6A vulnerability mapping in mouse DRG. (A) UMAP of 521 cells colored by Leiden-defined cell type. (B) Same UMAP colored by magnitude-weighted m6A loss burden (sites/kb). (C) m6A loss burden in sensory neurons versus non-neuronal cells. Welch’s t-test.

We projected the bulk m6A vulnerability data onto the single-cell landscape by mapping Cluster 3 (m6A erosion) genes to the single-cell gene expression matrix and computing a magnitude-weighted m6A loss burden per cell. This metric weights each cell not only by how many vulnerable genes it expresses but also by the magnitude of m6A loss at each gene, providing a biologically meaningful measure of epitranscriptomic exposure.

UMAP visualization revealed a striking spatial segregation of m6A loss burden (Figure 11B), with sensory neuron clusters (nociceptors and mechanoreceptors) showing substantially higher vulnerability scores than non-neuronal populations. Quantitative comparison (Figure 11C) confirmed this difference. Sensory neurons (n = 135) exhibited a mean m6A loss burden of 137,799 sites/kb compared to 62,712 sites/kb in non-neuronal cells (n = 386), representing a 2.2-fold enrichment. This difference was highly significant by both parametric (Welch’s t = 11.40, p = 4.98 × 10⁻²⁷) and non-parametric (Mann-Whitney p = 3.59 × 10⁻¹⁹) tests, with a large effect size (Cohen’s d = 1.14).

Cell-type-resolved analysis (Figure S6) showed that both nociceptors and mechanoreceptors exhibited similarly high m6A loss burden, with no significant difference between these two sensory neuron subtypes (p_adj = 0.73), indicating a shared vulnerability phenotype. All neuronal versus non-neuronal comparisons were significant (p_adj < 0.001 for all 10 pairwise tests), while comparisons among non-neuronal cell types were generally non-significant. The preceding analyses established that a transcript’s architectural features predetermine a transcript’s epitranscriptomic response to perturbation. Extending this principle from individual transcripts to cell populations generates a straightforward prediction: the cells with transcriptomes most enriched for high-m6A-density transcripts should bear the greatest aggregate modification loss. Without prior knowledge of the clinical phenotype, and using only transcript architecture and cell-type expression profiles, sensory neurons emerged as the most vulnerable population, computationally rediscovering the defining feature of BIPN. This validates the principle scales from individual molecules to cell-type-specific disease susceptibility.

## DISCUSSION

This study introduces the concept of architectural determinism: the principle that a transcript’s built-in structural features predetermine its epitranscriptomic response to perturbation. The m6A fate of a transcript under bortezomib treatment could be read from transcript-intrinsic features — motif composition and spatial organization — before any modification was measured or any drug exposure occurred. Standalone models using only these sequence-level features, with no m6A measurement, predicted 80–88% of the response variance. A multi-output machine learning model, trained on 69 measurable features of each transcript with no prior biological knowledge, returned a near-complete prediction of m6A response alongside the biological pathways most affected and the cell types most vulnerable. The model independently discovered known biology and generated testable mechanistic hypotheses from intrinsic RNA features alone, potentially establishing architectural determinism as both a biological principle and an analytical framework.

The architectural information encoded in transcripts carries genuine regulatory meaning. This is validated by the convergence of the computational predictions on the bortezomib–HIF1α–metabolic switch axis we have previously characterized through direct metabolic and protein-level measurements. Our published work established that bortezomib increases glycolysis while suppressing oxidative phosphorylation[20; 21]. The pipeline presented here, operating with no access to that prior knowledge, arrived at the same conclusion purely from transcript architecture. The dominant m6A erosion cluster enriched for precisely these metabolic pathways, and western blot data confirmed protein-level suppression of the same OXPHOS components. Critically, these transcripts would not have been identified by conventional differential expression analysis. The OXPHOS genes showed no significant changes in mRNA abundance. Their regulation was visible only at the epitranscriptomic and protein levels.

Architectural determinism accessed this biology precisely because it operates on a regulatory layer that standard transcriptomic workflows do not interrogate. This independent rediscovery of experimentally established biology from sequence features alone demonstrates that architectural encoding is not a statistical artifact but reflects a layer of regulation with measurable downstream consequences.

Architectural determinism also enabled a form of inference not available to conventional approaches. Transcript-level molecular vulnerabilities were mapped onto cell-type identity by integrating bulk-derived epitranscriptomic erosion data with single-nuclei transcriptomic profiles, identifying the cell types most affected. In this system, the analysis pointed to sensory neurons, consistent with the clinical presentation of BIPN[6; 31]. For any perturbation where architectural determinism operates, the same logic could potentially identify affected cell types, with implications for predicting drug toxicity targets and disease-vulnerable cell populations.

Independent human clinical evidence strengthens this interpretation. Polymorphisms in methylenetetrahydrofolate reductase (*Mthfr*, rs1801131, rs1801133, rs17421511) are associated with increased risk of BIPN in patients. These patients also exhibit lower *Mthfr* mRNA expression and elevated serum homocysteine[36]. *Mthfr* governs the production of S-adenosylmethionine (SAM)[10], the universal methyl donor required by the METTL3/METTL14 writer complex for m6A deposition. Individuals with reduced *Mthfr* activity have a constrained SAM supply and would therefore have impaired capacity to deposit or restore m6A marks after perturbation. The model identified the same vulnerability that clinical geneticists discovered independently, that impaired methylation capacity drives neuropathy risk, and potentially provides a mechanistic explanation for why these patients are susceptible.

Several limitations warrant consideration. The mechanism by which m6A erosion is induced remains to be determined, and whether it reflects reduced substrate availability, enhanced eraser activity, or altered co-transcriptional deposition, requires direct experimental investigation. How m6A erosion at specific transcripts leads to the observed changes in OXPHOS and glycolytic protein levels is also not yet established. The bidirectional remodeling at PABP recognition sites and the protein-level changes in the absence of mRNA abundance changes provided mechanistic clues, but confirming a direct link between m6A remodeling, PABP binding, and translational output will require targeted studies. The study used a mouse model, though the *Mthfr* pharmacogenomic data suggest conservation of the underlying mechanism in humans.

Architectural determinism was discovered here from a dataset comprising only three post-transcriptional regulatory layers and a single perturbation, yet the model achieved near-perfect predictive accuracy. Those same three layers are already implicated in a broad range of human diseases[2; 3; 7; 12; 14; 18; 23; 30; 38; 39]. Over 170 RNA modifications have been identified[17], the vast majority of which remain unstudied in the context of perturbation response. If three layers were sufficient to reveal deterministic encoding of this magnitude, the regulatory complexity embedded in the full epitranscriptomic landscape is likely far greater than currently appreciated. As the tools to measure these modifications on native RNA molecules continue to mature, the same analytical framework can be applied to uncover novel architectural vulnerabilities, which cell types bear the greatest burden, and through what mechanisms that vulnerability is realized. Ultimately, this knowledge could inform the design of targeted therapeutic strategies that anticipate and counteract epitranscriptomic disruption before it manifests as disease.

## Supporting information

Supplemental digital content

---

Figure S1. Scatter plots of predicted versus actual values on the held-out test set (n = 6,700 transcripts) for 15 perturbation-induced response variables. Each panel shows the model’s prediction accuracy for one response variable, with the dashed line representing perfect prediction. (A) expression_log2fc (R² = 0.377); (B) Δprob_modified (R² = 0.411); (C) Δmean_polyA (R² = 0.149); (D) Δmedian_polyA (R² = 0.121); (E) ΔpolyA_cv (R² = 0.169); (F) ΔpolyA_iqr (R² = 0.093); (G) ΔpolyA_mad (R² = 0.100); (H) ΔpolyA_skewness (R² = 0.142); (I) ΔpolyA_kurtosis (R² = 0.178); (J) Δpct_short_polyA (R² = 0.179); (K) Δpct_long_polyA (R² = 0.167); (L) polyA_cv_ratio (R² = 0.189); (M) polyA_mean_ratio (R² = 0.164); (N) Δisoform_entropy (R² = 0.351); (O) isoform_switch (R² = 0.286).

Figure S2. Top 20 features by importance for individual response variable models. (A) Expression fold-change model (expression_log2fc, R² = 0.377). (B) m6A modification probability model (Δprob_modified, R² = 0.411). (C) Isoform switching model (isoform_switch, R² = 0.286).

Figure S3. Focused validation of m6A erosion predictability. (A) Predicted versus actual Δsites/kb for the m6A erosion model. R² = 0.983; MAE = 343.96 Δsites/kb. Solid black line: linear fit; dashed gray line: perfect prediction. (B) Cross-validation fold stability across all five feature set configurations. All configurations achieve R² between 0.980 and 0.985 across all folds. (C) Distribution of model residuals (actual − predicted Δsites/kb). Residuals are centered near zero (mean = 0.817, std = 742.163 Δsites/kb), indicating no systematic prediction bias. (D) Permutation test. The observed R² = 0.983 (red line) vastly exceeds the null distribution (p value approaches zero). (E) Feature category ablation analysis showing mean R² (± s.d. across folds) as feature sets are progressively expanded from baseline m6A density only (6 features) to all features (51 features). (F) Feature category contribution to total Gini importance in the full model. m6A density features account for 99.5% of total importance. (G) Linear relationship between baseline m6A site density (Control, sites/kb) and perturbation-induced m6A loss (Δsites/kb). Pearson r = −0.992; regression slope = −0.735. (H) Temporal dynamics of m6A erosion at Day 7 (red) and Day 14 (blue). Steeper erosion at D7 (slope = −0.774) compared to D14 (slope = −0.696) indicates partial recovery during the post-treatment phase.

Figure S4. Isoform-resolved m6A motif-by-position heatmaps for each of the five *Ldha* isoforms expressed in DRGs. Δ m6A density (%) is shown by DRACH motif identity (y-axis) and normalized transcript position (x-axis, 0–100%) for D7 vs Control (upper) and D14 vs Control (lower). (A) *Ldha*-201, (B) *Ldha*-202, (C) *Ldha*-209, (D) *Ldha*-210, (D) *Ldha*-214.

Figure S5. Isoform-resolved for potential RBP target motif heatmaps for each of the five *Ldha* isoforms. Δ m6A density at RBP recognition motifs (y-axis) is shown as a function of normalized transcript position (x-axis) for D7 vs Control (upper) and D14 vs Control (lower). (A) *Ldha*-201, (B) *Ldha*-202, (C) *Ldha*-209, (D) *Ldha*-210, (E) *Ldha*-214.

Figure S6. Distribution of m6A loss burden across all seven DRG cell types shown as violin plots. Red lines: means; blue lines: medians. Pairwise Mann-Whitney U tests with FDR correction; statistical results in accompanying table.

Table 1. K-means clustering of 33,583 transcripts into four epitranscriptomic response patterns based on 15 normalized response variables. Cluster assignments, sizes, and mean response values for each regulatory layer are shown.

Table S1. Multi-output model response variables. Left: the 16 molecular response variables predicted by the gradient boosting model, organized by regulatory layer (m6A, poly(A), isoform, expression), with descriptions, units, computation methods, and individual R² values. Right: summary statistics by regulatory layer.

Table S2. Baseline transcript features used for model training. Left: the 69 baseline (control-condition) transcript features, organized by category (structural/motif, spatial, baseline m6A, baseline poly(A), baseline isoform, baseline expression), with descriptions, units, and data sources. Right: summary of feature categories.

Table S3. Gene set enrichment analysis for Cluster 3. Enrichr output for genes assigned to K-means Cluster 3, queried against GO Biological Process, GO Cellular Component, GO Molecular Function, KEGG, and Reactome libraries. Columns include term name, p-value, adjusted p-value, odds ratio, combined score, gene count, and gene list.

Table S4. Marker gene expression across single-nucleus RNA sequencing clusters. Percentage of nuclei expressing canonical marker genes within each of seven clusters (C0–C6) identified by unsupervised clustering of single-nucleus RNA sequencing data from mouse lumbar DRG (see Figure 10A). Markers are grouped by cell-type category: nociceptor, mechanoreceptor, glial, fibroblast, endothelial, and pan-neuronal. Cluster identities were assigned based on combinatorial expression patterns of established marker genes. Gene symbols follow standard mouse nomenclature.

## ACKNOWLEDGEMENTS

Supported by R01CA249939 and the Department of Neural and Pain Sciences. The authors declare there are no conflicts of interest.

## Code Availability

https://github.com/MelemedjianLab/drg-m6a-architectural-determinism

## Data sharing

Data are available upon reasonable request to the authors.

