## Supplemental digital content for "Transcript architecture predetermines m6A remodeling and sensory neuron vulnerability in chemotherapy-induced peripheral neuropathy"

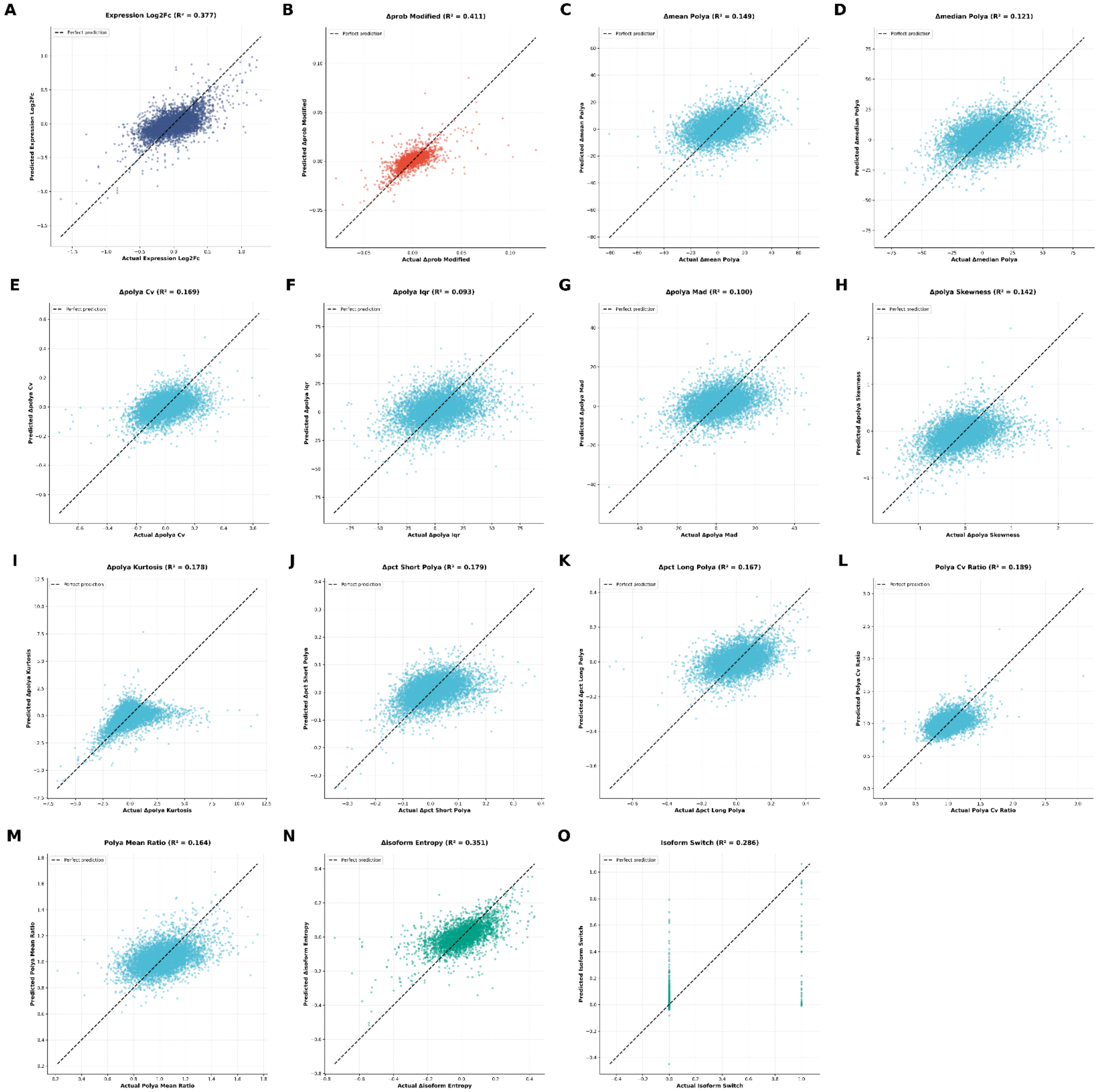


Figure S1. Scatter plots of predicted versus actual values on the held-out test set (n = 6,700 transcripts) for 15 perturbation-induced response variables. Each panel shows the model's prediction accuracy for one response variable, with the dashed line representing perfect prediction. (A) expression_log2fc ($R^2$ = 0.377); (B) Δprob_modified ($R^2$ = 0.411); (C) Δmean_polyA ($R^2$ = 0.149); (D) Δmedian_polyA ($R^2$ = 0.121); (E) ΔpolyA_cv ($R^2$ = 0.169); (F) ΔpolyA_iqr ($R^2$ = 0.093); (G) ΔpolyA_mad ($R^2$ = 0.100); (H) ΔpolyA_skewness ($R^2$ = 0.142); (I) ΔpolyA_kurtosis ($R^2$ = 0.178); (J) Δpct_short_polyA ($R^2$ = 0.179); (K) Δpct_long_polyA ($R^2$ = 0.167); (L) polyA_cv_ratio ($R^2$ = 0.189); (M) polyA_mean_ratio ($R^2$ = 0.164); (N) Δisoform_entropy ($R^2$ = 0.351); (O) isoform_switch ($R^2$ = 0.286).

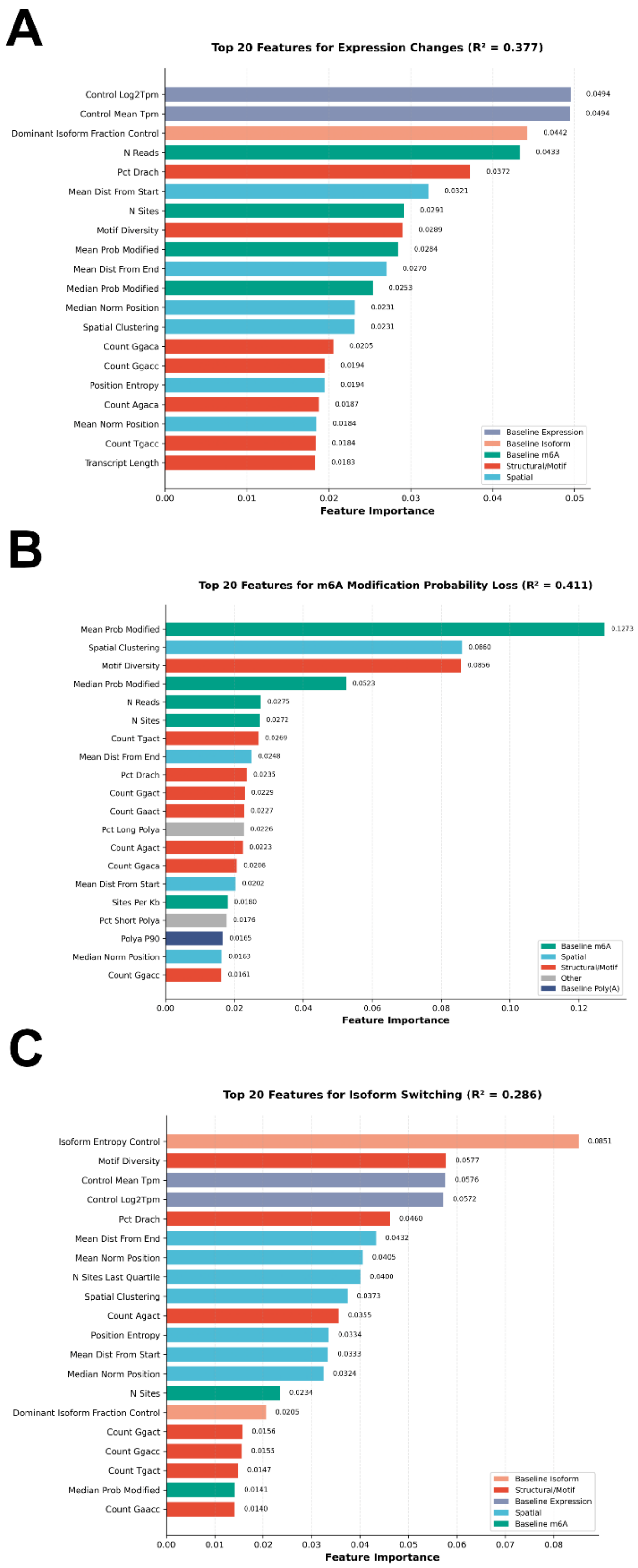


Figure S2. Top 20 features by importance for individual response variable models. (A) Expression fold-change model (expression_log2fc, $R^2 = 0.377$). (B) m6A modification probability model (Δprob_modified, $R^2 = 0.411$). (C) Isoform switching model (isoform_switch, $R^2 = 0.286$).

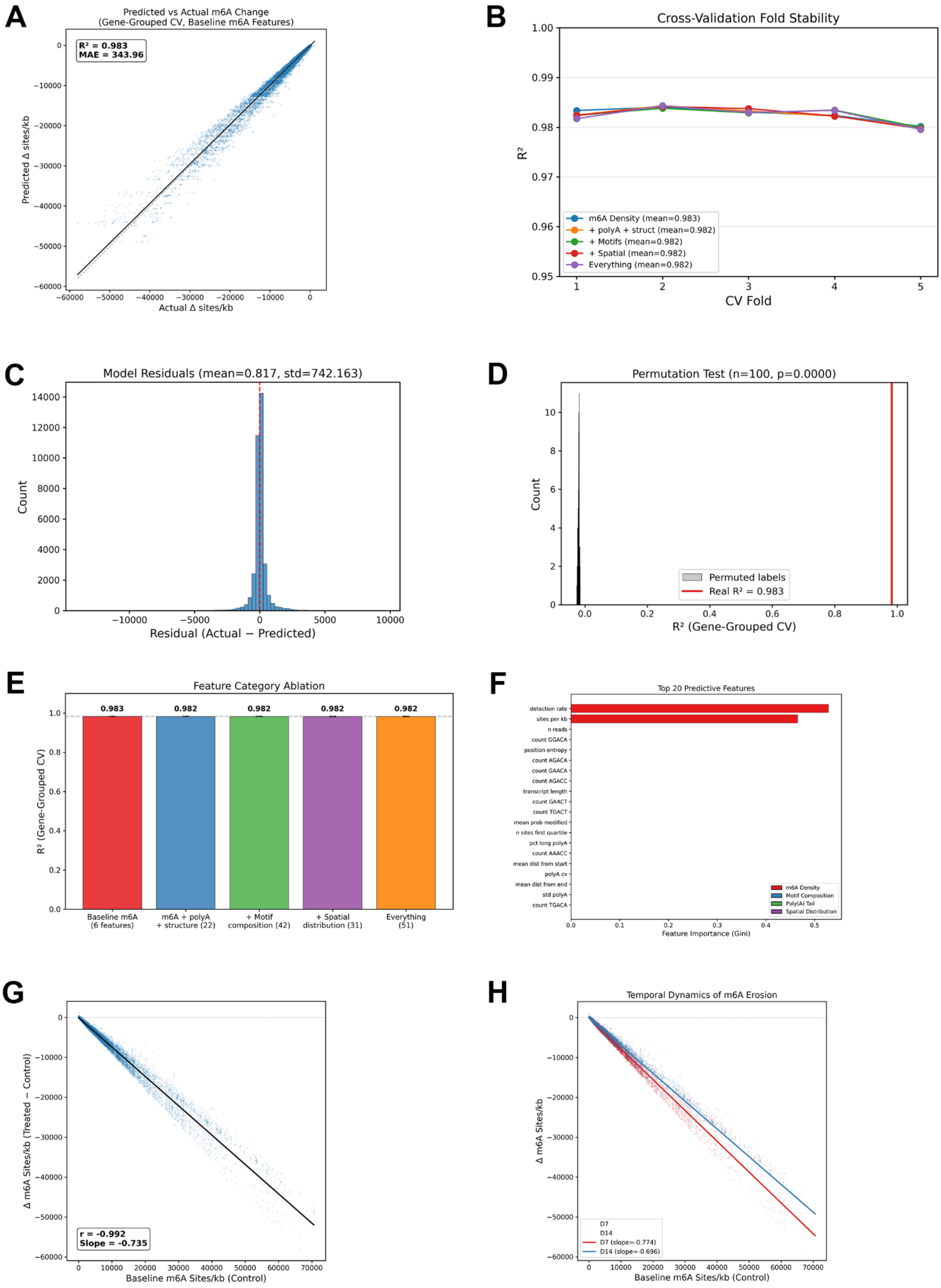

A
Predicted vs Actual m6A Change
(Gene-Grouped CV, Baseline m6A Features)
R² = 0.983
MAE = 343.96
Predicted Δ sites/kb
Actual Δ sites/kb
B
Cross-Validation Fold Stability
R²
CV Fold
m6A Density (mean=0.983)
+ polyA + struct (mean=0.982)
+ Motifs (mean=0.982)
+ Spatial (mean=0.982)
Everything (mean=0.982)
C
Model Residuals (mean=0.817, std=742.163)
Count
Residual (Actual − Predicted)
D
Permutation Test (n=100, p=0.0000)
Count
Permuted labels
Real R² = 0.983
R² (Gene-Grouped CV)
E
Feature Category Ablation
R² (Gene-Grouped CV)
0.983
0.982
0.982
0.982
0.982
Baseline m6A (6 features)
m6A + polyA + structure (22)
+ Motif composition (42)
+ Spatial distribution (31)
Everything (51)
F
Top 20 Predictive Features
detection rate
sites per kb
n reads
count GGACA
position entropy
count AGACA
count GAACA
count AGACC
transcript length
count GAACT
count TGACT
mean prob modified
n sites first quartile
pct long polyA
count AAACC
mean dist from start
polyA cv
mean dist from end
std polyA
count TGACA
m6A Density
Motif Composition
Poly(A) Tail
Spatial Distribution
Feature Importance (Gini)
G
Δ m6A Sites/kb (Treated − Control)
r = -0.992
Slope = -0.735
Baseline m6A Sites/kb (Control)
H
Temporal Dynamics of m6A Erosion
Δ m6A Sites/kb
D7
D14
D7 (slope=-0.774)
D14 (slope=-0.696)
Baseline m6A Sites/kb (Control)

Figure S3. Focused validation of m6A erosion predictability. (A) Predicted versus actual Δsites/kb for the m6A erosion model under gene-grouped 5-fold cross-validation using only 6 baseline m6A density features. $R^2$ = 0.983; MAE = 343.96 Δsites/kb. Solid black line: linear fit; dashed gray line: perfect prediction. (B) Cross-validation fold stability across all five feature set configurations. All configurations achieve $R^2$ between 0.980 and 0.985 across all folds. (C) Distribution of model residuals (actual − predicted Δsites/kb). Residuals are centered near zero (mean = 0.817, std = 742.163 Δsites/kb), indicating no systematic prediction bias. (D) Permutation test (n = 100 iterations with shuffled target labels). The observed $R^2$ = 0.983 (red line) vastly exceeds the null distribution (p value approaches zero). (E) Feature category ablation analysis showing mean $R^2$ (± s.d. across 5 CV folds) for models trained with progressively expanded feature sets: baseline m6A density only (6 features, $R^2$ = 0.983), m6A + poly(A) + structure (22 features, $R^2$ = 0.982), plus motif composition (42 features, $R^2$ = 0.982), plus spatial distribution (31 features, $R^2$ = 0.982), and all features (51 features, $R^2$ = 0.982). (F) Feature category contribution to total Gini importance in the full model (51 features). m6A density features account for 99.5% of total importance, with motif composition (0.3%), poly(A) tail (0.1%), and spatial distribution (<0.1%) contributing negligibly. (G) Linear relationship between baseline m6A site density (Control, sites/kb) and perturbation-induced m6A loss (Δsites/kb). Pearson r = −0.992; regression slope = −0.735. (H) Temporal dynamics of m6A erosion at Day 7 (peak pain, red) and Day 14 (recovery, blue). Steeper erosion at D7 (slope = −0.774) compared to D14 (slope = −0.696) indicates partial recovery during the post-treatment phase.

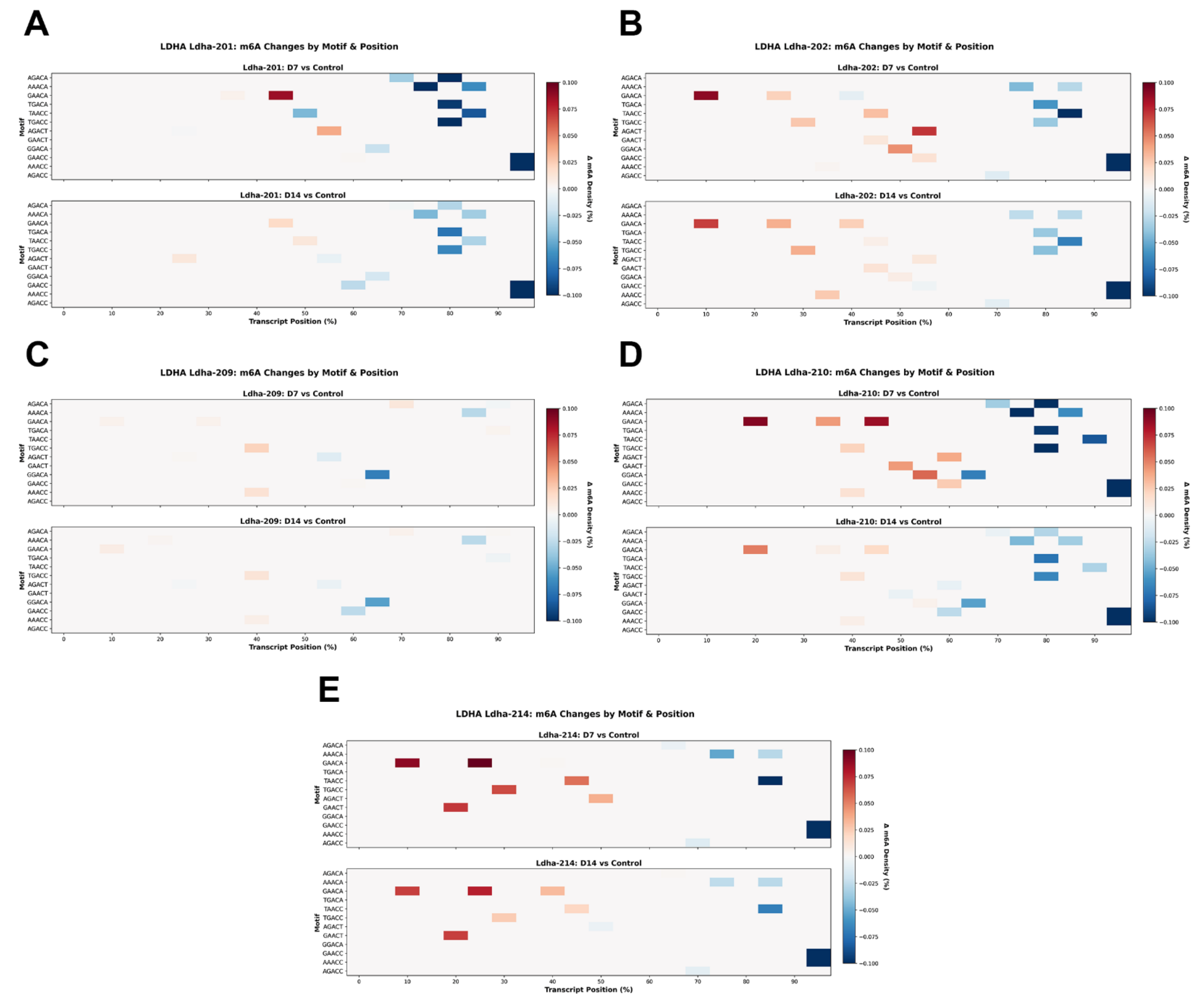


Figure S4. Isoform-resolved m6A motif-by-position heatmaps for each of the five LDHA isoforms expressed in DRGs. Δ m6A density (%) is shown by DRACH motif identity (y-axis) and normalized transcript position (x-axis, 0–100%) for D7 vs Control (upper) and D14 vs Control (lower). (A) Ldha-201, (B) Ldha-202, (C) Ldha-209, (D) Ldha-210, (D) Ldha-214.

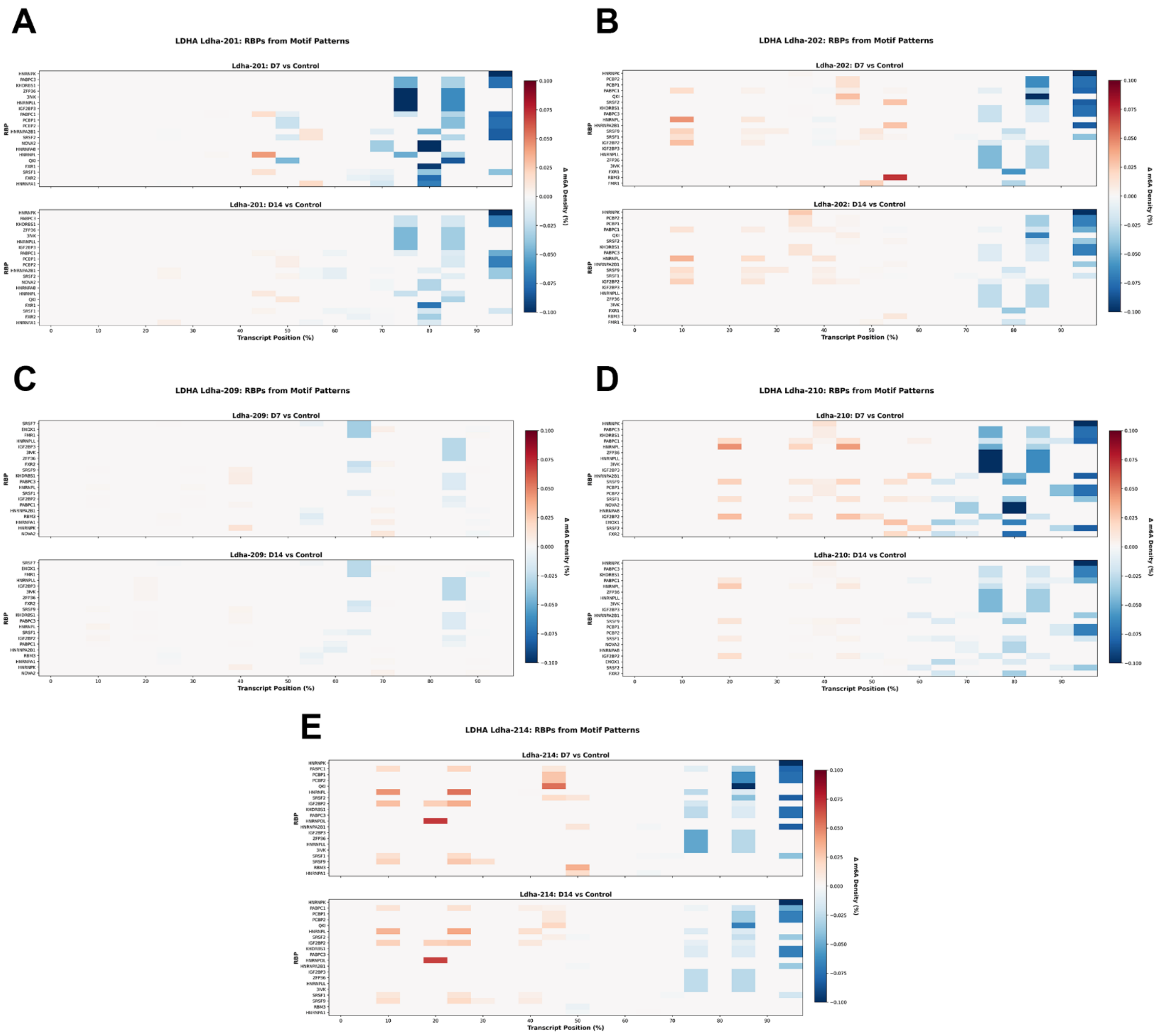


Figure S5. Isoform-resolved for potential RBP target motif heatmaps for each of the five LDHA isoforms. Δ m6A density at RBP recognition motifs (y-axis) is shown as a function of normalized transcript position (x-axis) for D7 vs Control (upper) and D14 vs Control (lower). (A) Ldha-201, (B) Ldha-202, (C) Ldha-209, (D) Ldha-210, (E) Ldha-214.

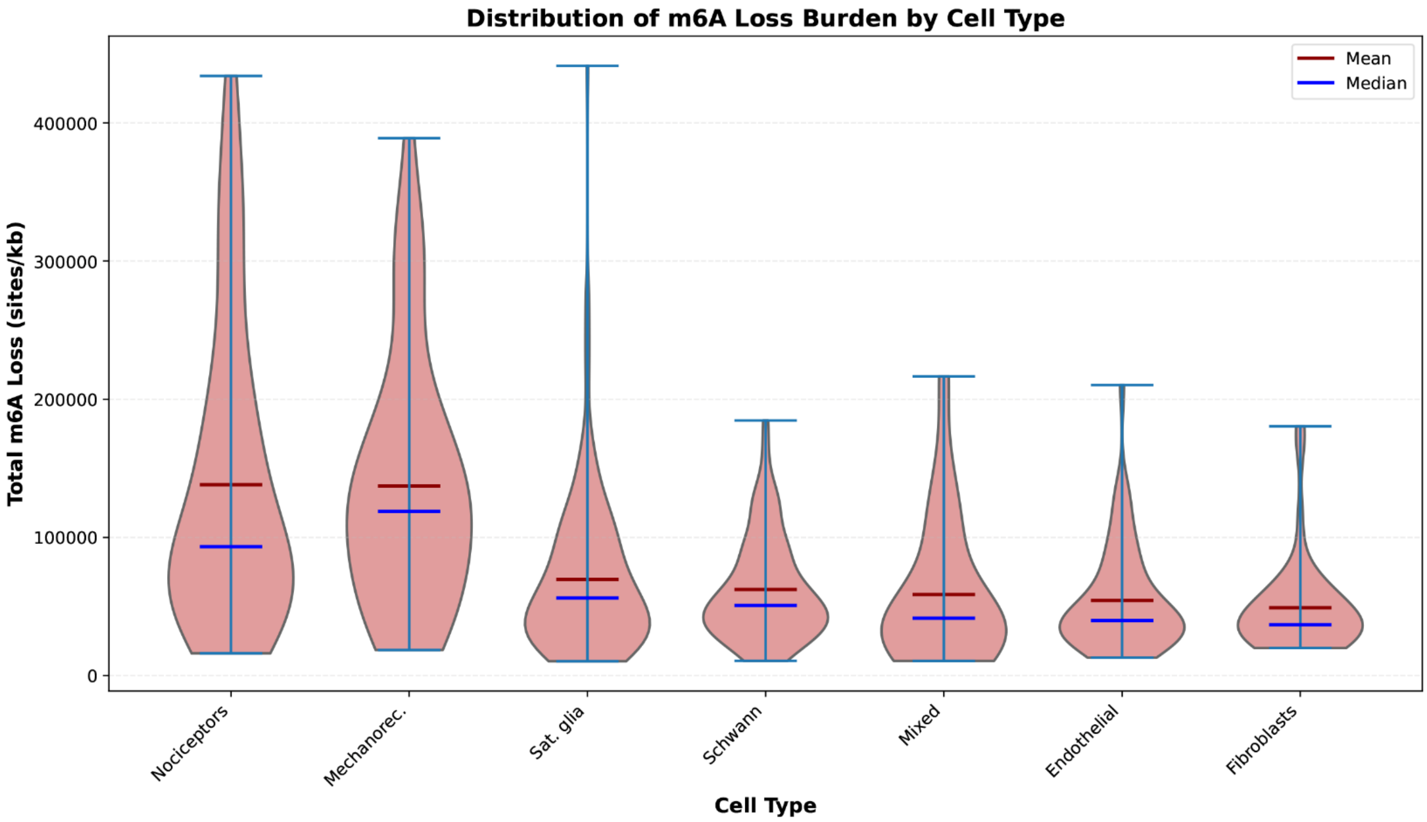


| Cell_Type_1 | Cell_Type_2 | U_statistic | p_value | mean_diff | p_adjusted | significant_raw | significant_fdr |
|---|---|---|---|---|---|---|---|
| Nociceptors | Fibroblasts | 2775 | 4.97E-09 | 89032.15258 | 6.94E-08 | TRUE | TRUE |
| Nociceptors | Schwann | 7667 | 6.61E-09 | 75811.91994 | 6.94E-08 | TRUE | TRUE |
| Mechanorec. | Schwann | 4600 | 1.86E-08 | 74811.39873 | 1.14E-07 | TRUE | TRUE |
| Nociceptors | Sat. glia | 9505 | 2.28E-08 | 68448.12012 | 1.14E-07 | TRUE | TRUE |
| Nociceptors | Endothelial | 3442 | 2.71E-08 | 83681.39888 | 1.14E-07 | TRUE | TRUE |
| Mechanorec. | Endothelial | 2034 | 6.45E-08 | 82680.87767 | 2.26E-07 | TRUE | TRUE |
| Mechanorec. | Fibroblasts | 1596 | 7.62E-08 | 88031.63137 | 2.29E-07 | TRUE | TRUE |
| Mechanorec. | Sat. glia | 5690 | 8.75E-08 | 67447.59891 | 2.30E-07 | TRUE | TRUE |
| Mechanorec. | Mixed | 823 | 7.90E-05 | 78496.55771 | 0.00018439 | TRUE | TRUE |
| Nociceptors | Mixed | 1397 | 0.000108454 | 79497.07892 | 0.000227753 | TRUE | TRUE |
| Schwann | Fibroblasts | 3003 | 0.010605726 | 13220.23264 | 0.020247296 | TRUE | TRUE |
| Sat. glia | Fibroblasts | 3631 | 0.04403322 | 20584.03246 | 0.077058136 | TRUE | FALSE |
| Schwann | Endothelial | 3649 | 0.059086546 | 7869.478933 | 0.095447498 | FALSE | FALSE |
| Sat. glia | Endothelial | 4460 | 0.146999706 | 15233.27875 | 0.206595665 | FALSE | FALSE |
| Schwann | Mixed | 1523 | 0.147568332 | 3685.158975 | 0.206595665 | FALSE | FALSE |
| Sat. glia | Mixed | 1896 | 0.20098113 | 11048.9588 | 0.263787732 | FALSE | FALSE |
| Endothelial | Fibroblasts | 1081 | 0.483773337 | 5350.753707 | 0.597602357 | FALSE | FALSE |
| Nociceptors | Mechanorec. | 1999 | 0.622785598 | 1000.52121 | 0.726583198 | FALSE | FALSE |
| Mixed | Endothelial | 510 | 0.756771064 | 4184.319958 | 0.836431175 | FALSE | FALSE |
| Mixed | Fibroblasts | 396 | 0.84032502 | 9535.073665 | 0.882341271 | FALSE | FALSE |
| Sat. glia | Schwann | 9346 | 0.965275916 | 7363.79982 | 0.965275916 | FALSE | FALSE |

Figure S6. Distribution of m6A loss burden across all seven DRG cell types shown as violin plots. Red lines: means; blue lines: medians. Pairwise Mann-Whitney U tests with FDR correction; statistical results in accompanying table.
